# An autocrine Vitamin D-driven Th1 shutdown program can be exploited for COVID-19

**DOI:** 10.1101/2020.07.18.210161

**Authors:** Reuben McGregor, Daniel Chauss, Tilo Freiwald, Bingyu Yan, Luopin Wang, Estefania Nova-Lamperti, Zonghao Zhang, Heather Teague, Erin E West, Jack Bibby, Audrey Kelly, Amna Malik, Alexandra F Freeman, Daniella Schwartz, Didier Portilla, Susan John, Paul Lavender, Michail S Lionakis, Nehal N Mehta, Claudia Kemper, Nichola Cooper, Giovanna Lombardi, Arian Laurence, Majid Kazemian, Behdad Afzali

**Affiliations:** Immunoregulation Section, Kidney Diseases Branch, National Institute of Diabetes and Digestive and Kidney Diseases (NIDDK), NIH, Bethesda, MD, USA; Department of Molecular Medicine and Pathology, School of Medical Sciences, The University of Auckland, Auckland, New Zealand; Departments of Biochemistry and Computer Science, Purdue University, West Lafayette, IN, USA; Molecular and Translational Immunology Laboratory, Department of Clinical Biochemistry and Immunology, Faculty of Pharmacy; Universidad de Concepcion, Concepcion, Chile; Department of Agricultural and Biological Engineering, Purdue University, West Lafayette IN, USA; Laboratory of Inflammation & Cardiometabolic diseases, Cardiovascular Branch, National Heart, Lung, and Blood Institute (NHLBI), National Institutes of Health (NIH), Bethesda, MD, USA; Laboratory of Molecular Immunology and the Immunology Center, National Heart, Lung, and Blood Institute (NHLBI), National Institutes of Health (NIH), Bethesda, MD, USA; School of Immunology and Microbial Sciences, Faculty of Life Sciences and Medicine, King’s College London, London, UK; Department of Medicine, Imperial College London, London, UK; Laboratory of Clinical Immunology & Microbiology, National Institute of Allergy and Infectious Diseases (NIAID), NIH, Bethesda, MD, USA; Laboratory of Allergic Diseases, National Institute of Allergy and Infectious Diseases (NIAID), NIH, Bethesda, MD, USA; Division of Nephrology and the Center for Immunity, Inflammation and Regenerative Medicine, University of Virginia, VA, USA; Fungal Pathogenesis Section, Laboratory of Clinical Immunology and Microbiology, National Institute of Allergy and Infectious Diseases (NIAID), NIH, Bethesda, MD, USA; Institute for Systemic Inflammation Research, University of Lübeck, Lübeck, Germany; MRC Centre for Transplantation, School of Immunology and Microbial Sciences, Faculty of Life Sciences and Medicine, King’s College London, London, UK; Nuffield Department of Medicine, University of Oxford, UK

**Keywords:** SARS-CoV2, COVID-19, Complement, Vitamin D, single cell RNA-sequencing, STAT3, BACH2, c-JUN

## Abstract

Pro-inflammatory immune responses are necessary for effective pathogen clearance, but cause severe tissue damage if not shut down in a timely manner^1,2^. Excessive complement and IFN-γ-associated responses are known drivers of immunopathogenesis^3^ and are among the most highly induced immune programs in hyper-inflammatory SARS-CoV2 lung infection^4^. The molecular mechanisms that govern orderly shutdown and retraction of these responses remain poorly understood. Here, we show that complement triggers contraction of IFN-γ producing CD4^+^ T helper (Th) 1 cell responses by inducing expression of the vitamin D (VitD) receptor (VDR) and CYP27B1, the enzyme that activates VitD, permitting T cells to both activate and respond to VitD. VitD then initiates the transition from pro-inflammatory IFN-γ^+^ Th1 cells to suppressive IL-10^+^ Th1 cells. This process is primed by dynamic changes in the epigenetic landscape of CD4^+^ T cells, generating superenhancers and recruiting c-JUN and BACH2, a key immunoregulatory transcription factor^5–7^. Accordingly, cells in psoriatic skin treated with VitD increased BACH2 expression, and BACH2 haplo-insufficient CD4^+^ T cells were defective in IL-10 production. As proof-of-concept, we show that CD4^+^ T cells in the bronchoalveolar lavage fluid (BALF) of patients with COVID-19 are Th1-skewed and that VDR is among the top regulators of genes induced by SARS-CoV2. Importantly, genes normally down-regulated by VitD were de-repressed in CD4^+^ BALF T cells of COVID-19, indicating that the VitD-driven shutdown program is impaired in this setting. The active metabolite of VitD, alfacalcidol, and cortico-steroids were among the top predicted pharmaceuticals that could normalize SARS-CoV2 induced genes. These data indicate that adjunct therapy with VitD in the context of other immunomodulatory drugs may be a beneficial strategy to dampen hyperinflammation in severe COVID-19.

---

Severe acute respiratory syndrome coronavirus (SARS-CoV)-2, causing coronavirus disease 2019 (COVID-19), is a novel virus responsible for the largest pandemic of the 21^st^ century. A significant number of patients with COVID-19 develop severe and life-threatening acute respiratory distress syndrome, characterized by hyper-inflammation and respiratory failure^8^. Mortality from severe COVID-19 remains high, due in part to limited understanding of the pathophysiology of the disease and the limited range of specific immunomodulatory therapies available. Indeed, the efficacy of dexamethasone in reducing mortality in COVID-19 suggests the importance of inflammation to disease severity^9^. Survivors of COVID-19, and those with milder disease, may lose significant normal tissue functions due to persistent inflammation and fibrosis^10^, leading to the development of chronic lung disease^11^. To gain insights into the pathophysiologic mechanisms of COVID-19, we analyzed single cell RNAseq (scRNAseq) data from the bronchoalveolar lavage fluid (BALF) and the peripheral blood mononuclear cells (PBMCs) of patients with COVID-19 and healthy controls. Protective immunity to both SARS-CoV1 and MERS-CoV, two closely related coronaviruses, is thought to be mediated by, amongst other cells, IFN-γ producing airway memory CD4^+^ T helper (Th) cells^12^. Development of Th1-polarized responses is also a feature of SARS-CoV2 in humans^13^. Moreover, severe COVID-19 is accompanied by prolonged and exacerbated Th1 responses^14^, which are suspected to contribute towards the observed pathogenic hyper-inflammation. We therefore focused our analyses of the scRNAseq datasets on CD4^+^ T cells. We identified the T cell populations within scRNAseq of BALF (**Figs. S1a-b**) and re-analyzed them to uncover 7 main subclusters, including two CD4^+^ Th cell clusters according to the well-characterized markers of these cells (**Figs. 1a and S1c**). Although the proportion of CD4^+^ cells within T cells represented in samples from patients and controls did not differ (**Fig. 1a**), 312 genes were upregulated and 134 genes down-regulated in patient CD4^+^ T cells compared to controls (**Fig. 1b** and **Table S1a**). Functional annotation of these differentially expressed genes (DEGs) showed enrichment for noteworthy pathways, including interferon-γ response and complement (**Fig. 1c** and **Table S1b**). Indeed, BALF CD4^+^ T cells in patients with COVID-19 appeared to be skewed towards the Th1 as opposed to the Th2 or Th17 lineages (**Figs 1d-e**). Similar examination within scRNAseq of PBMC from patients with COVID-19 and healthy controls (**Figs. S2a-b**) did not show meaningful differences in expression of Th1, Th2 or Th17 lineage genes (**Fig. S2c**), suggesting that expression of the Th1 program is a specific feature of Th cells at the site of pulmonary inflammation where virus-specific T cells may be concentrated^13^.

**Fig 1.**
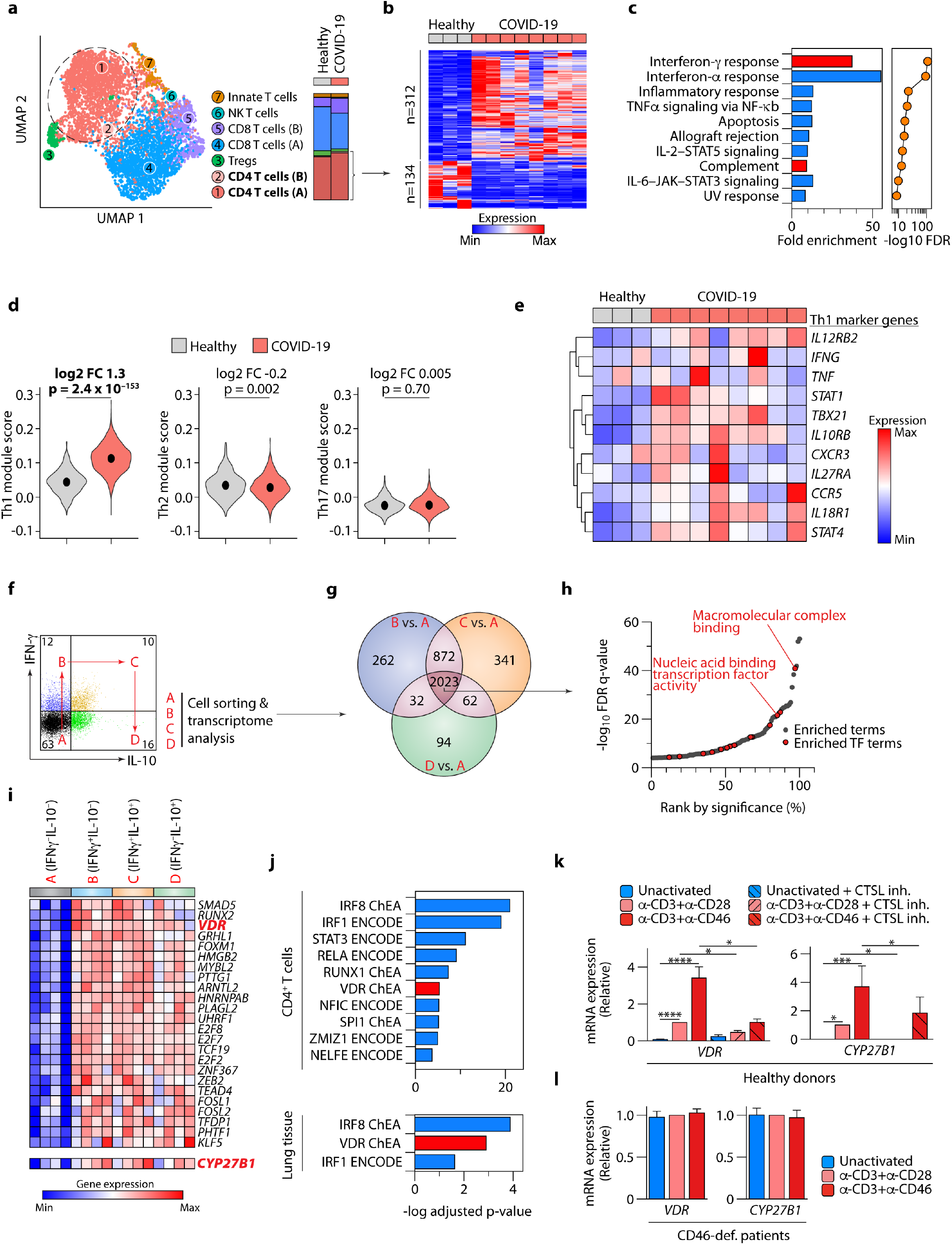
*VDR* and *CYP27B1* are induced by complement and predicted to be regulators of the Th1 program in COVID-19. **a**, UMAP projection of scRNAseq showing sub-clustering of T cells from (*n*=8) bronchoalveolar lavage fluid (BALF) of patients with COVID-19 and (*n*=3) healthy controls. Stack bars (right) show cumulative cellularities across samples in patients and controls. Dot plot of marker genes for these clusters are shown in **Fig. S1c**. Highlighted are clusters 1 and 2, representing CD4^+^ T cells. **b-c**, Heatmap showing differentially expressed genes (DEGs; at least 1.5 fold-change in either direction at adjusted p-value <0.05) between CD4^+^ T cells of patients with COVID-19 and healthy controls (**b**) and enrichment of those DEGs in Hallmark MSigDB genesets (**c**). Highlighted in red in **c** are hallmark interferon-γ response and complement pathways. **d**, Violin plots showing expressions of Th1, Th2 and Th17 specific genes, respectively, summarized as module scores, in BALF CD4^+^ T cells of patients with COVID-19 and healthy controls. Medians are indicated. **e**, Heatmap showing mean expression of classic Th1 marker genes in BALF CD4^+^ T cells of patients with COVID-19 and healthy controls. **f**, Representative flow cytometry plot showing IFN-γ and IL-10 expression in CD4^+^ T cells activated with anti(α)-CD3+α-CD46 and the 4 quadrants (marked as A, B, C and D) from which cells were FACS-sorted for transcriptome analysis. Live and single cells are pre-gated. **g**, Venn diagram showing numbers of DEGs (1.5-fold change in either direction at p-value<0.05) comparing cells in quadrants B, C and D against A, respectively (*n*=4 experiments). **h**, Enrichment of gene ontology molecular function terms in shared DEGs (intersect of Venn diagram in **g**) of α-CD3+α-CD46-activated CD4^+^ T cells, ranked by statistical significance. Marked in red are terms corresponding to transcription factor (TF) activity. **i**, Heatmap showing induced TFs in α-CD3+α-CD46-activated CD4^+^ T cells at each stage of the Th1 life-cycle shown in **f**. Highlighted in red is the vitamin D receptor (*VDR*). Also shown is expression of *CYP27B1* (highlighted in red). **j**, EnrichR-predicted ENCODE and ChEA (ChIP enrichment analysis) TFs regulating the DEGs between COVID-19 *vs*. healthy donor CD4^+^ BALF T cells (upper panel) and lung biopsies (lower panel). **k**, Expression of *VDR* (left panel) and *CYP27B1* (right panel) mRNA in CD4^+^ T cells activated, or not, as indicated, in the presence of absence of a cell-permeable cathepsin L inhibitor (CTSL inh.) (*n*=5 experiments). **l**, Expression of *VDR* (left panel) and *CYP27B1* (right panel) mRNA in CD4^+^ T cells of a patient with CD46 deficiency, activated, or not, as indicated (*n*=3). Data in **a-d** and **upper panel of j** are sourced from GSE145926 and GSE122960. Data in **lower panel** of **j** are from GSE147507. Data in **f-i** are from GSE119416. Data in **l** are from microarrays in ^67^. Bars in **k-l** show mean + sem; *p<0.05, **p<0.01, ***p<0.001, ****p<0.0001 by ANOVA with Kruskal-Wallis test. Exact p-values in **d** have been calculated using two-tailed Wilcoxon tests.

Enrichment of the complement system in local T cells (**Fig. 1c**) was notable because a) we have recently identified complement as one of the most highly induced pathways in CD4^+^ T cells infiltrating into lung tissues^15^, b) SARS-CoV2 is a potent inducer of complement, especially complement factor 3 (C3), in lung epithelial cells^4^ and c) others have previously identified the lungs of patients with COVID-19 as a complement-rich environment^16^. C3 is cleaved to generate activation fragments, of which C3b binds CD46 receptors on T cells. We have shown that CD4^+^ T lymphocytes in the lungs of patients with COVID-19 have a CD46-activated signature^4^. CD46, engaged by T cell-generated C3b in an autocrine fashion, works co-operatively with T cell receptor stimulation to drive both Th1 differentiation and subsequent T cell shutdown^17^. Thus, T cells activated *ex vivo* with anti(α)-CD3 and α-CD46 produce IFN-γ, then co-produce IL-10 before shutting down IFN-γ to produce only IL-10^17^ (**Fig. 1f**) (please note that this CD46 system is specific to human CD4^+^ T cells and is not present in mouse T cells). The switch from effector T cells, important for pathogen clearance, into IL-10 producing cells reduces collateral damage and is a natural transition in a T cell’s life-cycle^18^. This suggests that IL-10 is produced by cells that are successfully transitioning into the Th1 shut program^17^. Indeed, in models of *T. gondii* and *T. cruzi* infections, mice unable to produce IL-10 clear infection more rapidly but die of severe tissue damage from uncontrolled Th1 responses^1,2^. We asked whether prolonged Th1 responses in COVID-19 patients are due to the failure in initiating this Th1 shutdown program. *IL10* mRNA, similar to most cytokines, was not detectable in the scRNAseq data, but detectable in bulk RNAseq from BALF. Consistent with the scRNA-seq data, bulk RNAseq from BALF demonstrated a significant enrichment of Th1-related genes in transcripts more highly expressed in cells from patients with COVID-19 compared to control – but a ~4-fold reduction in *IL10* levels (**Fig. S1d**), consistent with the notion that Th1 cells indeed did not initiate the shutdown program.

To examine how complement normally regulates the Th1 cell life-cycle and particularly their shutdown program, we next studied Th cells of healthy subjects. Unless specified otherwise, we used Treg-depleted CD4^+^ Th cells (CD4^+^CD25^-^) throughout. After activation with α-CD3+α-CD46, we FACS-sorted Th cells from each quadrant of the Th1 life-cycle, by surface cytokine capture (**Figs. 1f** and **S3a-c**) and performed transcriptome analysis. Comparing the transcriptomes of IFN-γ^+^IL-10^-^, IFN-γ^+^IL-10^+^ and IFN-γ^-^IL-10^+^ against IFN-γ^-^IL-10^-^ Th cells revealed approximately 2000 DEGs in common (**Figs. 1g** and **S1d** and **Tables S1c-d**). These genes were significantly enriched for factors whose molecular function pertained to transcription factor (TF) biology (**Figs. S3e-f** and **1h** and **Table S1e**), indicating that regulation of TFs is key to CD46 biology. In total, 24 TFs were induced by CD46 (**Fig. 1i**), among which was the vitamin D receptor (VDR) (**Fig. 1i**). Our attention was drawn to VDR for two reasons. First, because independent prediction of the top TFs regulating the DEGs of CD4^+^ BALF T cells and lung biopsies from patient versus healthy donors returned VDR among the top candidates (**Fig. 1j** and **Table S1f**). Second, we noted that *CYP27B1* was concurrently induced in the transcriptome data (**Fig. 1i**). CYP27B1 is the vitamin D (VitD) 1α-hydroxylase enzyme, responsible for the final activation step of VitD, catalyzing the conversion of 25(OH)-VitD to biologically active 1,25(OH)_2_-VitD^19^. Inducible expression of *CYP27B1* and *VDR* in Th cells indicated the likely presence of an autocrine/paracrine loop, whereby T cells can both activate and respond to VitD. Although both have been described as induced in activated T cells^20,21^, the molecular mechanism and functional consequences are currently unknown. α-CD3+α-CD28 stimulation activates intracellular C3 processing by cathepsin L (CTSL) in T cells, which generates autocrine C3b to ligate CD46 on the cell surface^22^. CD46 itself enhances intracellular C3 processing to generate more C3b^22^. To establish that both *CYP27B1* and *VDR* are transcriptionally induced by complement, we confirmed that both α-CD3+α-CD28 and α-CD3+α-CD46 induced these genes in T cells and that this effect was nullified by addition of a cell soluble CTSL inhibitor to the cultures (**Fig. 1k**). Similarly, T cells from CD46 deficient patients were unable to upregulate either gene with either stimulus *in vitro* (**Fig. 1l**). Collectively, these data indicated that complement components ligating CD46 on the surface of Th cells induce both the enzyme and the receptor to be able to fully activate and also to respond to VitD.

VitD is a steroid hormone with pleiotropic functions in the immune system, including both anti-microbial as well as regulatory properties, which are cell and context-dependent^23^. VitD deficiency is associated with higher prevalence and adverse outcomes from infectious diseases, especially respiratory infections, such as influenza, tuberculosis and viral upper respiratory tract illnesses^24,25^, as well as several autoimmune diseases, including type 1 diabetes mellitus, multiple sclerosis, rheumatoid arthritis and inflammatory bowel disease^26^. Th cells play key roles in all these diseases. Thus, understanding the consequences of VDR ligation in Th cells is potentially of importance for many diseases including COVID-19. For all subsequent experiments, unless specified otherwise, Th cells were activated with α-CD3 and α-CD28, and we used the active form of VitD (1α,25(OH)_2_-VitD). We confirmed that VDR protein is induced by T cell activation (**Fig. S4a**), addition of active VitD enhances VDR expression^27^ (**Fig. S4a**), and VitD-bound VDR translocates to the nucleus (**Figs. S4b-c**). Th cells cultured with VitD upregulated 296 and down-regulated 157 genes (**Figs. S4d** and **2ab** and **Tables S2a-b**), which were not attributable to changes in cell proliferation or death (**Fig. S4e**). We noted that classical VitD-regulated genes, including *CTLA4*^28^, *CD38*^29^ and *CYP24A1*^30^, were induced by VitD and that both type 1 (*IFNG*) and type 3 (*IL17A, IL17F, IL22* and *IL26*) cytokines were repressed (**Figs 2a-b**), as has been previously reported^28^. The genes induced by VitD included two cytokines, *IL10* and *IL6*, as well as several TFs including *JUN, BACH2* and *STAT3* (**Figs 2a-b**). A population of potently suppressive IL-10 producing CD4^+^FoxP3^-^ cells, termed type 1 regulatory T cells^31^ can be induced from naïve Th cells in response to VitD^32^. In our data, genes induced/repressed in VitD-treated cells were not enriched in the gene signature of Tr1 cells (**Fig. S4f**) and did not become CD49b^+^LAG3^+^, the archetypal surface phenotype of Tr1 cells^33^ (**Fig. S4g**). Similarly, we did not observe upregulation of *FOXP3*, the master TF of natural Tregs, by VitD treatment (**Table S2b**). The genes regulated by VitD were most significantly enriched for cytokines (**Fig. S4h**). Unexpectedly, in the hierarchy of cytokines regulated by VitD, *IL6*, normally a pro-inflammatory cytokine, was the most highly induced (**Fig. S5a**). We confirmed in supernatants of VitD or carrier treated Th cells that VitD repressed IFN-γ and IL-17 and induced IL-10 and IL-6 (**Fig. S5b**). Moreover, there was a dose-response relationship between VitD concentrations and these effects on Th cells (**Fig. 2c**). To confirm that activation-induced expression of *CYP27B1* (**Fig. S5c**) allows Th cells to activate and respond to VitD in an autocrine/paracrine manner, we provided inactive 25(OH)-VitD to Th cells and measured the same cytokines. We observed the same pattern as before – repression of IFN-γ and IL-17 and induction of IL-10 and IL-6, indicating that the T cells had converted 25(OH)-VitD to active 1,25(OH)_2_-VitD themselves (**Fig. S5d**).

**Fig. 2.**
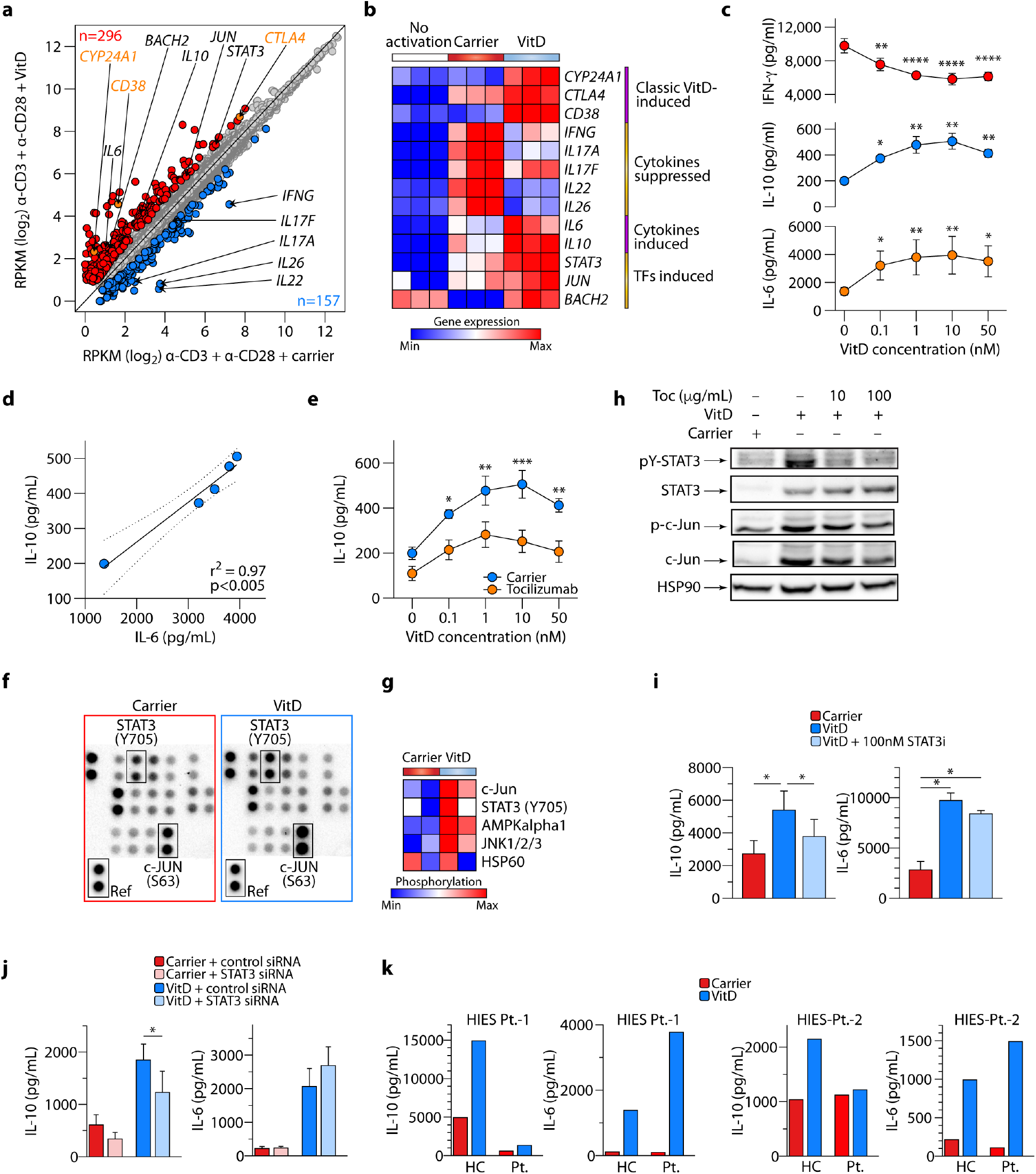
Vitamin D induces IL-10 in CD4^+^ T cells by IL-6-STAT3 signal transduction. **a**, Scatter plot showing mRNA expression (RPKM) of all genes in CD4^+^ T cells treated with VitD or carrier. VitD-induced and -repressed genes (at least 1.75-fold change in either direction at FDR<0.05) are depicted in red and blue, respectively. Noteworthy genes are annotated (black), including classical VitD-induced genes (orange). *n*=3 independent experiments. **b**, Heatmap showing expression of select genes in CD4^+^ T cells treated with VitD or carrier. **c**, concentrations of indicated cytokines in supernatants of CD4^+^ T cells treated for 72h with increasing concentrations of VitD. Stars indicate statistically significant changes in comparison to 0nM of VitD. **d**, Pearson correlation between concentrations of IL-6 and IL-10 in culture supernatants of CD4^+^ T cells treated with VitD. Shown is the correlation line, plus 95% confidence interval. **e**, Concentrations of IL-10 in supernatants of CD4^+^ T cells cultured with increasing concentrations of VitD, with and without Tocilizumab. **f**, Representative image from *n*=2 experiments of a phospho-kinase array (array of 43 kinases in duplicate spots) carried out on 3-day lysates of CD4^+^ T cells treated with carrier or VitD. Location of STAT3 phosphorylated at lysine 705, c-JUN phosphorylated at serine 63 and reference spot (to which all spots are normalized) are indicated. **g**, heatmap showing normalized phosphorylation values of differentially phosphorylated proteins following VitD-treatment (please see also **Fig. S6a**) in *n*=2 donors. **h**, Western blots of lysates of CD4^+^ T cells treated with carrier or VitD with, and without, Tocilizumab (Toc) at the concentrations shown. Shown are representative images of pY-STAT3, STAT3, p-c-JUN, c-JUN, with Hsp90 as loading control, from *n*=3 experiments (quantified in **Fig S6b**). **i-j**, IL-10 and IL-6 production from CD4^+^ T cells cultured with carrier or VitD, with or without a STAT3 inhibitor (STAT3i) (**i**) or transfected with control siRNA or siRNA targeting STAT3 (**j**). **k**, Production of IL-10 and IL-6 from CD4^+^ T cells of *n*=2 healthy controls (HC) or patients (Pt.) with Hyper IgE syndrome (HIES Pt.) treated with carrier or VitD. Unless indicated, all cells in **Fig. 2** have been activated with α-CD3+α-CD28. Cumulative data in **c**, **e** and **i-j** (all n=3 independent experiments) depict mean+sem. *p<0.05, **p<0.01, ***p<0.001, ****p<0.0001 by one-way (**c**, **i**, **j**) and two-way ANOVA (**e**).

IL-6 is a pleiotropic cytokine, with almost all stromal and all cells of the immune system being capable of producing it^34^, but not previously linked with induction by VitD in T cells. We therefore established that *IL6* mRNA and protein were produced by T cells and induced by VitD (**Fig. S5e**) and that VitD treatment significantly increased the percentage of Th cells positive for intracellular IL-6 (**Fig. S5f**). In these experiments, it was striking that there appeared to be a strong correlation between IL-6 and IL-10 concentrations produced in response to VitD (**Fig. 2d**), suggesting that one of these cytokines may drive expression of the other. Accordingly, we stimulated Th cells with VitD in the presence or absence of Tocilizumab, an IL-6 receptor (IL-6R) blocking antibody used clinically for the management of IL-6-dependent cytokine release syndrome, including the suggested cytokine storm in COVID-19. Tocilizumab significantly impaired the production of IL-10 in response to VitD, indicating that IL-6R signaling induced by autocrine/paracrine IL-6 induces IL-10 in Th cells (**Fig. 2e**). IL-6 can co-operate with IL-27^35^ or IL-21 ^36^ to promote IL-10 production from mouse T cells. However, both IL-21 and IL-27 were repressed by VitD in our transcriptomes (**Table S2b**), so it does not appear that these cytokines are co-operating with IL-6 to induce IL-10 in this setting. Addition of IL-6, without VitD (although T cells can activate VitD, the ‘substrate’ still needs to be provided in cultures for conversion), to Th cultures did not induce IL-10 (**Fig. S5g**) but increased pro-inflammatory IL-17 (**Fig S5h**), as has been reported^37,38^. These data indicate that the pro-inflammatory functions of IL-6 may be restricted or averted by the production of antiinflammatory IL-10 in the presence of VitD in human Th cells (**Fig. 2e**). A study of VitD supplementation in Mongolian children noted significant increases in serum IL-6 (as well as a non-significant increase in IL-10)^39^ indicating that our observations can also occur *in vivo*. In the skin, the site of the highest levels of VitD, overexpression of IL-6 provides protection from injurious stimuli or infection^40^ and IL-6 deficiency impairs cutaneous wound healing^41^. Moreover, one of the adverse effects of anti-IL-6R for the treatment of inflammatory arthritis has been idiosyncratic development of psoriasis^42,43^, indicating a tolerogenic role for IL-6 at this site.

Genes differentially expressed by VitD treatment were enriched in cytokine signaling pathways (**Fig. S4h**), which are commonly mediated by phosphoproteins. To further delineate the molecular mechanisms underlying the immunoregulatory properties of VitD, we therefore carried out a phospho-kinase protein array using lysates of carrier and VitD-treated cells. Five proteins showed significant differences in phosphorylation between carrier and VitD, most notably c-JUN and STAT3, both of which were significantly more phosphorylated following treatment with VitD (**Figs. 2f-g** and **S6a**). By Western blotting we confirmed that STAT3 protein and STAT3 phosphorylation were both induced by VitD (**Fig. 2h** and **S6b**). IL-6 is a potent signal driving STAT3 activation by phosphorylation^34^. Accordingly, STAT3 phosphorylation, but not protein expression induced by VitD, was abrogated by blockade of the IL-6R with Tocilizumab, indicating that IL-6 induced by VitD is responsible for the phosphorylation but not the expression of STAT3 protein (**Fig. 2h** and **S6b**). By contrast, c-JUN protein expression and c-JUN phosphorylation were both dependent on VitD and mostly independent of IL-6 (**Fig. 2h** and **S6b**). Since VitD-induced STAT3 phosphorylation was mediated by IL-6, we investigated whether STAT3 is the driver of IL-10 produced by VitD. Both culture of VitD-treated Th cells with a cell permeable STAT3 inhibitor and knock-down of STAT3 by siRNA significantly impaired IL-10 produced in response to VitD (**Figs. 2i-j**). Likewise, Th cells from two rare patients with hyper-IgE syndrome that have dominant negative mutations in *STAT3*, failed to produce significant IL-10 in response to VitD (**Fig. 2k**). Collectively, these data established that VitD induces STAT3 and IL-6, and IL-6 phosphorylates STAT3 to promote production of IL-10.

VitD-bound VDR interacts with histone acetyl transferases^44^, transcriptional co-activators^45^, co-repressors^46^, and chromatin remodeling complexes^47^ to modulate transcription. We therefore explored the effect of VitD on the epigenetic landscape of T cells, using FACS-sorted memory CD4^+^ T cells as these cells express the VDR without requiring pre-activation. We carried out cleavage under target and release using nuclease (CUT&RUN)^48^ for histone 3 lysine 27 acetylation (H3K27Ac), a marker of active regions of the genome, in VitD and carrier-treated primary CD4^+^ T cells. VitD treatment induced dynamic changes in histone acetylation genome-wide over time, as shown in **Figs. 3a-b**, indicating that enhancer formation induced by VDR is a common mechanism by which VitD affects gene expression. As expected, loci containing genes whose transcription was increased by VitD showed increased histone acetylation and loci with genes whose transcription was repressed had reduced histone acetylation (**Fig. S7a**). By 48h after VitD treatment, there were approximately 25000 induced H3K27Ac peaks and approximately 21000 repressed peaks (**Fig. 3c** and **Table S3a**). The size of these peaks was significantly affected by VitD addition, leading to the generation of new super-enhancer (SE) architectures and promotion of existing SEs (**Fig. 3d**). SEs are complex regulatory domains comprised of collections of enhancers that are critical for regulating genes of particular importance to cell identity and risk of genetic disease^7,49,50^. VitD-modified SEs included those associated with *BACH2, STAT3, IL10* and other genes induced by VitD (**Fig. 3d** and **Table S3b**). To identify potential TFs recruited to these loci, we carried out TF DNA motif finding at H3K27Ac peak loci induced by VitD. Native CUT&RUN, as well as ChIP-seq for VDR, proved challenging in primary Th cells; however, the top enriched motifs in the H3K27Ac peaks induced by VitD were VDR, AP-1 family members, notably c-JUN and BACH2, and STAT3 (**Fig. 3e**), all three of which were TFs whose transcription was also induced by VitD (**Figs. 2a-b**). c-JUN is an AP-1 basic leucine zipper (bZIP) family member, whose primary function is DNA transcription^51^. Since we had also noted activating phosphorylation of c-JUN in response to VitD (**Figs. 2f-h**), we reasoned that c-JUN is likely to play an important role in regulation of genes by VitD. We carried out c-JUN CUT&RUN in VitD and carrier treated primary CD4^+^ T cells (**Fig. S7b**) and found significant recruitment of c-JUN to VitD-induced H3K27Ac loci over time, which superseded that induced by carrier treatment alone (**Figs. 3f** and **S7c** and **Table S3c**). In particular, we found c-JUN binding at loci containing genes of interest, including *CTLA4*, *STAT3* and *BACH2* (**Figs. 3g**, **S7d-e** and **2a**).

**Fig. 3.**
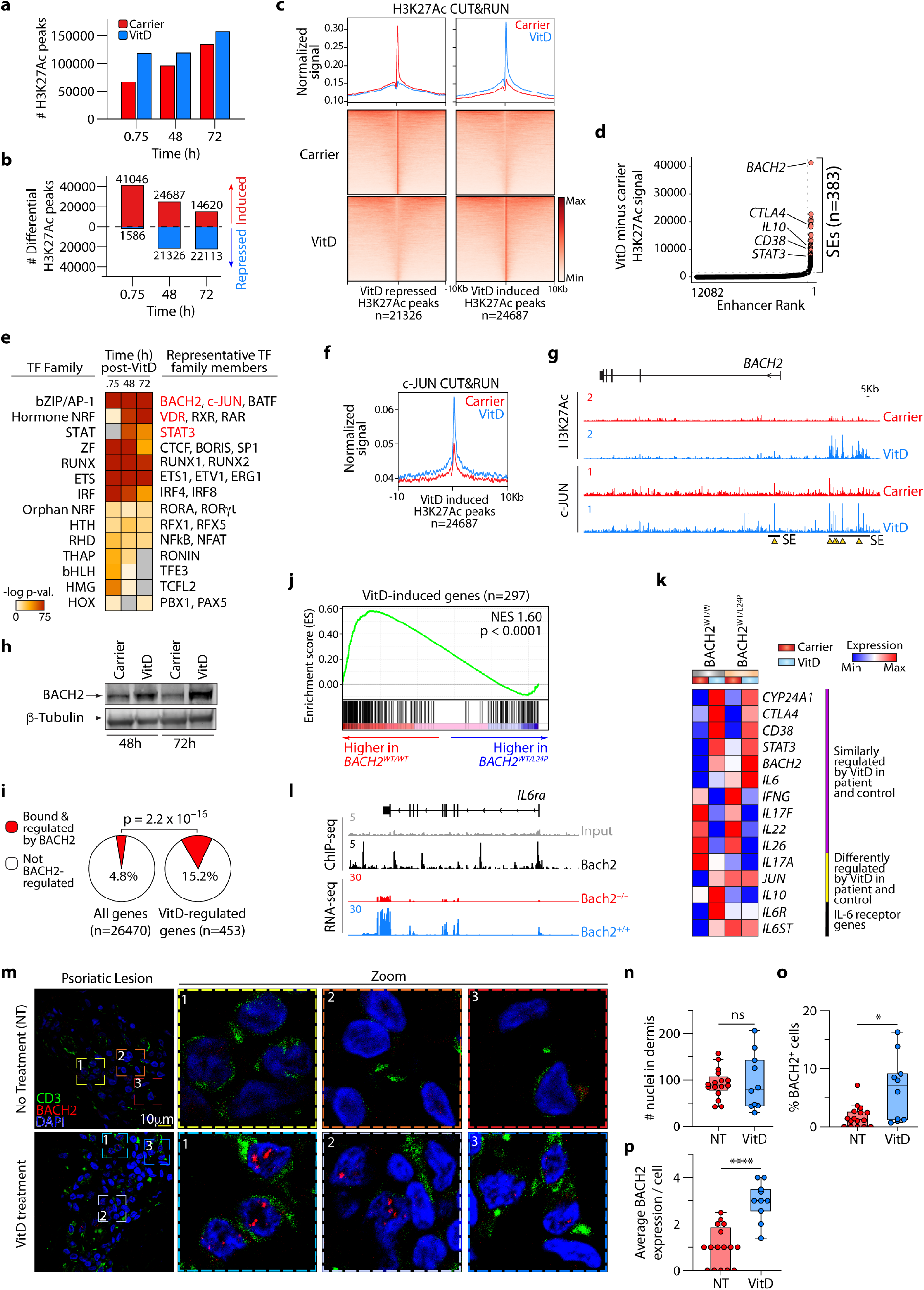
Vitamin D reshapes epigenetic landscape of CD4 T cells via transcription factors c-JUN and BACH2. **a**, Genome-wide H3K27Ac CUT&RUN peaks 45mins, 48h and 72h after VitD or carrier-treatment of CD4^+^ T cells. **b**, differential H3K27Ac peaks (signal ≥0.2, ≥1.5 fold change) after VitD or carrier-treatment of CD4^+^ T cells at the indicated time points. **c**, Heatmaps showing H3K27Ac signal at VitD repressed and VitD-induced peaks (below) and histograms showing normalized signals in carrier and VitD treated cells (48hr) above. **d**, Enriched transcription factor (TF) DNA motifs at H3K27Ac peak loci induced by VitD. Shown are TF families on the left and representative TF members enriched in the data on the right. **e**, Ranked order of H3K27Ac-loaded enhancers induced by VitD in Th cells after 48h. Super-enhancers (SEs) are indicated. Marked are the relative positions, ranked according to signal intensity (higher = greater signal intensity), of enhancers attributed to selected genes. **f**, Histograms showing normalized c-JUN CUT&RUN signal in VitD and carrier-treated CD4^+^ T cells (48 hrs), centered at VitD-induced H3K27Ac peaks. **g**, Representative genome browser tracks showing H3K27Ac and c-JUN CUT&RUN at the *BACH2* locus following VitD or carrier treatment of CD4^+^ T cells (48hr). Indicated is the position of the *BACH2* SE induced by VitD and select peaks of c-JUN and/or H3K27Ac peaks (yellow triangles). **h**, Immunoblot for BACH2 and β-Tubulin in lysates of VitD and carrier-treated CD4^+^ T cells at the indicated time points. Shown is a representative example from *n*=3 experiments. **i**, Pie-charts comparing the percentage of VitD-regulated genes that are bound and regulated by BACH2 against all genes in the genome. The Fisher exact p-value is shown. **j**, GSEA showing enrichment in VitD-induced genes when comparing the transcriptomes of VitD-treated healthy wild-type controls (BACH2^WT/WT^) with CD4^+^ T cells from a patient with haploinsufficiency of BACH2 (BACH2^WT/L24P^). NES = normalized enrichment score. **k**, Heatmap showing patterns of expression of select VitD-regulated genes in the transcriptomes of carrier and VitD-treated CD4^+^ T cells from a haplo-insufficient BACH2^WT/L24P^ patient and healthy haplo-sufficient BACH2^WT/WT^ control. **l**, genome browser tracks showing Bach2 ChIP-seq at the *IL6ra* locus and expression of *IL6ra* mRNA in CD4^+^ T cells of Bach2 wild-type (*Bach2^+/+^*) and knock-out (*Bach2*^-/-^) mice. Source data are indicated. **m-p**, Representative images of the dermis of lesional skin from psoriasis patients with and without VitD supplementation stained for BACH2 (red) and CD3 (green), showing overview (leftmost) and zoomed images (right three images) (**m**), the number of nuclei in each image (**n**), the percentage of BACH2 positive cells relative to nuclei frequency per image (**o**) and the average number of BACH2 foci per cell (**p**). Data in **m-p** are from *n*=3 patients without and *n*=2 patients with VitD treatment and are shown as mean + sem. Unless indicated, all *in vitro* T cell experiments depicted in **Fig. 3** have been activated with α-CD3+α-CD28. Data in **l** are from GSE45975. *p<0.01, ****p<0.0001 by unpaired two-tailed t-test.

BACH2 is a critical immunoregulatory TF^6,52^. Both haploinsufficiency and single nucleotide variants of this TF are associated with monogenic and polygenic autoimmune diseases of humans, respectively^7,53,54^. We confirmed that VitD induces BACH2 protein expression in Th cells (**Fig. 3h**) – most likely via promoting its super enhancer and c-JUN binding (**Figs. 3d** and **3g**) – and found that the genes regulated by VitD were approximately 3-fold enriched for those regulated by BACH2 (**Fig. 3i**). Geneset enrichment analysis (GSEA)^55^ further revealed that genes more highly expressed in VitD-treated cells are enriched in BACH2-induced genes and genes more highly expressed in carrier treated cells are enriched in BACH2-repressed genes (**Figs. S8a-c**). Among the most highly enriched in the leading edge of BACH2-induced genes was the IL-6 receptor alpha chain (*IL6R*) (**Figs. S8a-c**). These observations suggested that a significant portion of VitD driven transcription is BACH2 dependent. To confirm *in vivo* association between BACH2 and VitD, we took two parallel approaches: First, we compared the transcriptomes of VitD treated Th cells from healthy control (*BACH2^WT/WT^*) to those from a rare patient with a loss of function mutation in *BACH2 (BACH2^WT/L24P^*) resulting in haploinsufficiency^7^. VitD-induced genes were significantly enriched in the transcriptomes of control wildtype cells (**Fig. 3j** and **Table S4a**), indicating that normal concentrations of BACH2 are required for appropriate regulation of VitD-induced genes. We did not find the same pattern of enrichment for BACH2-repressed genes (**Fig. S9d**), potentially indicating that half the normal concentration of BACH2 protein is sufficient for repression of VitD targets. As a result, type 1 and type 3 inflammatory cytokines were normally repressed by VitD in the context of haploinsufficiency of BACH2 (**Fig. 3k** and **Table S4b**). Despite higher basal mRNA levels, *IL6* and *STAT3* were both induced by VitD when BACH2 levels were sub-normal, but *IL10* was not, signifying that the IL-6–STAT3–IL-10 axis is disrupted in the context of BACH2 haploinsufficiency (**Figs. S8e** and **3k**). Indeed, we found that *IL6R* is not appropriately induced when BACH2 levels are sub-normal (**Figs. S8e** and **3k**). Bach2 ChIP-seq and RNAseq from wild-type and *Bach2*^-/-^ animals confirmed that the *IL6r* gene is a direct genomic target of Bach2 and that knockout status for this TF significantly impairs the expression of *Il6r* (**Fig. 3l**). Second, we performed confocal imaging on the dermis of patients with psoriasis treated, or not, with VitD, because the psoriatic skin is rich in CD4^+^ T cells and is frequently treated successfully with VitD^56^. Indeed, VitD-treated psoriatic skin demonstrated significantly increased numbers of BACH2^+^ cells and greater numbers of intranuclear foci of BACH2^57^, compared to untreated skin (**Figs. 3m-p**), indicating that BACH2 induction by VitD occurs in an *in vivo* setting. Collectively, these data indicate that BACH2 is a VitD-induced protein that regulates a significant portion of the VitD transcriptome, with a key event being the induction of IL-6R that enables autocrine/paracrine IL-6-STAT3 signaling to induce IL-10.

There is compelling epidemiological association between COVID-19 incidence/severity and VitD deficiency/insufficiency, including higher mortality in patients with pigmented skin (of African or Indian descent)^58–63^. However, if causal, the molecular mechanisms are lacking. Given the apparent failure of normal Th1 contraction observed in COVID-19 patients (**Figs. 1d** and **S1d**), we hypothesized that VitD immunoregulation could be mechanistically important for the hyper-inflammation seen in the lungs of COVID-19 patients and may provide a rationale for therapy. To explore this possibility, we looked at expression of genes regulated by VitD (**Fig. 2a** and **Table S2b**) in BALF CD4^+^ T cells. Expression of VitD-repressed genes, summarized as the module score, was significantly higher in BALF Th cells of patients with COVID-19 than healthy controls (**Fig. 4a**). This was further corroborated by GSEA showing that genes expressed more highly in BALF Th cells of COVID-19 compared to healthy control were enriched in VitD-repressed genes (**Fig. 4b**). We noted that on a per cell basis the VitD-repressed module score correlated strongly with the Th1 score in BALF Th cells (**Fig. 4c**), indicating a reciprocal relationship between Th1 genes and VitD-repressed genes. In contrast, expression of VitD-induced genes was not substantially different between the two groups (**Fig. S9**), nor were VitD-regulated genes different between CD4^+^ T cells from PBMC of patients compared to healthy donors (**Fig. S10**). We constructed receiver operating characteristic (ROC) curves of the module scores for the Th1 and VitD-repressed gene signatures to assess the ability of the module scores to distinguish patient from healthy BALF Th cells on a per cell basis. The area under the curve (AUC) for the two scores were 0.91 and 0.80, respectively (p<0.00001 for both) (**Fig. 4d**), indicating that both Th1 and VitD-repressed gene signatures are strong features of COVID-19 CD4^+^ BALF T cells. To contextualize these performances, we additionally calculated the AUC for the module scores of every geneset in hallmark and canonical pathways curated by MSigDB (*n*=2279 genesets)^55,64,65^. The top performing sets were type I and type II interferon responses, as expected. The VitD-repressed geneset was within the top 1% of all genesets at distinguishing patient from healthy Th cells (**Fig. 4e** and **Table S5**), indicating that this is among the very best performing genesets. We next used EnrichR to predict 461 significant drugs that could potentially be used to counteract the genes upregulated in COVID-19 BALF Th cells. Five of the top 10 most significant drugs were steroids and two were VitD (alfacalcidol) (**Fig. 4f** and **Table S6**). Dexamethasone, a cortico-steroid drug, has recently been shown to reduce mortality from severe COVID-19^9^, presumably through immunosuppression. A significant number of the genes that were modulated by VitD were shared by cortico-steroids (**Fig. S11a**). Steroids, including dexamethasone, enhance the effects of VitD by increasing transcription of VDR^66^. These data suggest that, in patients with severe COVID-19, addition of VitD to other immunomodulatory agents might be beneficial. From experience in other diseases, it is unlikely that VitD will be effective as monotherapy. Combination therapy (at lower doses) could ameliorate the significant adverse effects of high dose steroid treatments, including over-immunosuppression or metabolic side-effects. In addition, an important consideration of VitD therapy in COVID-19 is the stimulation of IL-6 VitD from CD4^+^ T cells. Although IL-6 induces IL-10 in Th cells, it is a potential concern that IL-6 acting on other cells might have pro-inflammatory properties. These potential effects can be mitigated by adding VitD as an adjunct therapy to other immunomodulatory drugs, such as steroids or JAK inhibitors^4^.

**Fig. 4.**
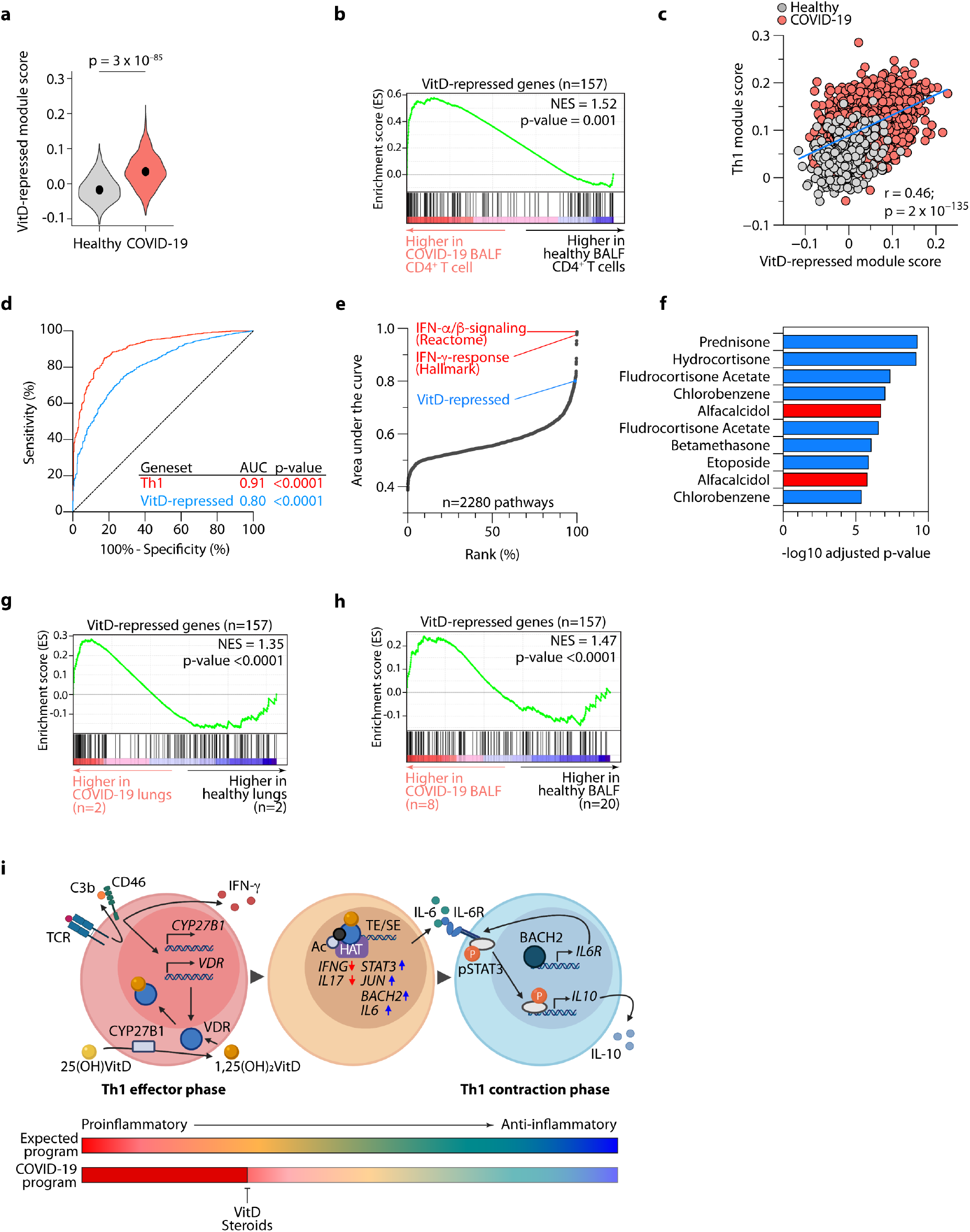
The exacerbated Th1 program in COVID-19 lungs can be retracted by Vitamin D. **a**, Violin plots showing expressions of VitD-repressed genes, summarized as module scores, in BALF CD4^+^ T cells of patients with COVID-19 and healthy controls. Exact p-values in **a** have been calculated using twotailed Wilcoxon tests. **b**, GSEA showing enrichment in VitD-repressed genes within genes more highly expressed in scRNAseq CD4^+^ BALF T cells of patients with COVID-19 compared to healthy controls. **c**, Correlation between module scores of Th1-genes and VitD-repressed genes on a per cell basis in BALF CD4^+^ T cells of patients with COVID-19 and healthy controls. Pearson r and exact p values are shown. **d**, Receiver operating characteristic (ROC) curve, evaluating the performance of the Th1 and VitD-repressed module scores to distinguish BALF CD4^+^ T cells of patients with COVID-19 from healthy controls. Shown are the area under the curve (AUC) statistics and p-values. **e**, Analyses showing the performance of all MSigDB canonical and hallmark genesets to distinguish BALF CD4^+^ T cells of patients with COVID-19 from healthy controls, ranked by AUC values. Marked are the top 2 performing genesets in red and the position of the VitD-repressed geneset within the top 1% of all genesets. **f**, Top 10 drugs predicted (out of 461 significant drugs) to counteract genes induced in BALF CD4^+^ T cells of COVID-19 patients compared to healthy controls, ordered by adjusted p-value. Highlighted in red is alfacalcidol, an FDA-approved active form of VitD. **g-h**, GSEA showing enrichment in VitD-repressed genes for genes more highly expressed in bulk RNA-seq lung biopsy specimens (**g**) and bulk RNA-seq BALF cells (**h**) of COVID-19 compared to healthy controls. NES = normalized expression value. **i**, Schematic model of autocrine VitD-driven Th1 contraction program and potential intervention of impaired COVID-19 program with VitD and cortico-steroids. Data in **a-f** are from *n*=8 patients with COVID-19 and *n*=3 healthy controls, sourced from GSE145926 and GSE122960. Data in **g-h** are from GSE147507 and HRA000143, respectively, and *n* numbers are indicated.

In summary, we have shown that complement initiates Th1 shutdown via orchestrating an autocrine/paracrine autoregulatory VitD loop. VitD causes epigenetic re-modelling, induces and recruits a set of TFs including STAT3, c-JUN and BACH2, that collectively repress Th1 and Th17 programs and induces IL-10 via IL-6-STAT3 signaling. We show that this program is down-regulated in the lungs of patients with severe COVID-19 and could be potentially exploited therapeutically by using adjunct VitD treatment in patients with COVID-19 (**Fig. 4i**).

## Acknowledgments

The authors thank patients who contributed samples toward this study. We would like to acknowledge Gennady Denisov and the NIH HPC (Biowulf) for their efforts in maintaining essential bioinformatic programs. We would like to thank Qian Zhu, Nan Liu, Derek Janssens, and Steven Henikoff for critical discussions related to CUT&RUN experimentation and data analysis. This work was supported by the Wellcome Trust (grant 097261/Z/11/Z to B.A), the Crohn’s and Colitis Foundation of America (grant CCFA no. 3765 — CCFA genetics initiative to A.L.), British Heart Foundation (grant RG/13/12/30395 to G.L.), National Institute of General Medical Sciences (R35GM138283 to M.K.), the Showalter Trust (research award to M.K.) and the National Agency of Research and Development of Chile (grant PAI79170073 to ENL). Research was also supported by the National Institute for Health Research (NIHR) Biomedical Research Centre based at Guy’s and St Thomas’ NHS Foundation Trust and King’s College London and/or the NIHR Clinical Research Facility. The views expressed are those of the author(s) and not necessarily those of the NHS, the NIHR or the Department of Health. This research was supported (in part) by the Intramural Research Programs of the National Institute of Diabetes and Digestive and Kidney Diseases (project number ZIA/DK075149 to B.A), the National Heart, Lung, and Blood Institute (project numbers ZIA/Hl006223 to C.K. and ZIA/HL006193 to N.M.) and the National Institute of Allergy and Infectious Diseases (project number ZIA/AI001175 to M.S.L) of the National Institutes of Health.

## Author Contributions

Single cell data analysis was performed by T.F., B.Y., Z.Z. and J.B. Bulk RNA experiments and analyses were carried out by T.F, D.C., L.W. and N.C. CUT & RUN and analysis was carried out by D.C., L.W., P.L. and A.K. Other wet lab experiments were conducted by R.M. E.N-L., H.T., E.E.W., C.K. B.A. and S.J. Patient samples were provided by D.S., N.M and N.C., who also analysed data. D.P, S.J., P.L., M.S.L., N.M., C.K., N.C., G.L., and A.L. provided intellectual input, interpreted data, provided supervision of wet lab work and helped write the paper. R.M., T.F., D.C., M.S.L., A.L., M.K. and B.A. wrote the manuscript. M.K. and B.A. conceived the project.

## Methods

### Ethics approvals

Human studies were conducted in accordance with the Declaration of Helsinki and approved by the Institutional Review Board of Guy’s Hospital (reference 09/H0707/86), National Institutes of Health (approval numbers 7458, PACI, 13-H-0065 and 00-I-0159) and Imperial College London (approval number 12/WA/0196 ICHTB HTA license number 12275 to project R14098). All patients provided informed written consent.

### Human T cell isolation and culture

Human PBMCs were purified from either anonymized leukodepletion cones (Blood Transfusion Service, NHS Blood and Transplantation) or healthy volunteer whole blood packs/buffy coats from the NIH Blood Bank and from fresh blood of patients. Leukodepletion cones were diluted 1 in 4 with sterile PBS and layered onto 15mL of lymphoprep (Axis-Shield). Whole blood packs/buffy coats were diluted 1 in 2 with sterile PBS onto 15mL Lymphocyte Separation Medium (25-072-CV LSM, Corning). Cells were separated by centrifugation at 20C for 20 min, with slow acceleration and no brakes and PBMC collected by harvesting interface cells.

CD4^+^CD25^-^ cells were used throughout the paper, unless specified below. RosetteSep Human CD4^+^ enrichment cocktail (Stem Cell) was used to enrich for purified CD4+ T cells from leukodepletion cones for both culture and pre-enrichment prior to sorting, as per manufacturers protocol. Briefly, cones where first diluted 1 in 2 and incubated at room temperature for 20 min with Rosettesep cocktail. Blood was then diluted a further 1 in 4 and CD4^+^ Tcells separated as described for PBMCs. CD4^+^CD25^-^ cells were then obtained by depletion of CD25^+^ T cells using CD25 positive selection (CD25 microbeads II, Human, Miltenyi Biotec) and collecting the unlabeled cells.

Human memory CD4^+^ T cells were used for RNA-seq and CUT&RUN experiments as these cells express the VDR, while naïve T cells require pre-activation first. Human memory CD4^+^ T cells were isolated either from PBMCs using Miltenyi Memory CD4^+^ T cell Isolation Kit human (130091893) or StemCell EasySep Human Memory CD4^+^ T Cell Enrichment Kit (19157) according to the manufacturers’ instructions or by FACS sorting, as detailed below.

Cells were cultured in Lonza X-VIVO-15 Serum-free Hematopoietic Cell Medium (04-418Q) supplemented with 50 IU/mL penicillin, 50μg/mL streptomycin (Invitrogen), 2mM L-glutamine (PAA Labarotories) at 37C 5% CO2. CD4^+^ memory T cells were cultured in a 96 well U bottom plate at a cell density of 10^6^ cells per mL in a total volume of 200μL. CD4^+^ cells were activated using Gibco Dynabeads Human T-Activator CD3/CD28 for T Cell Expansion and Activation (11131D) at a cell to bead ratio of 4:1. Where indicated, cells were activated in 96 well plates coated with antibodies against CD3 (OKT3, made inhouse by the Washington University hybridoma facility), CD3 + CD28 (CD28.2, Becton Dickinson) or CD3 + CD46 (TRA-2-10, a gift from John P Atkinson, Washington University, USA), all at 2μg/mL in sterile PBS overnight at 4C, in the presence and absence of a cell soluble cathepsin L inhibitor (ALX-260-133-M001 from Enzo Life Sciences, Exeter, UK) at 1μM final concentration. Cells were additionally cultured in the presence or absence of 1α,25-Dihydroxyvitamin D3 (Enzo Life Sciences) or 25-Dihydroxyvitamin D3 (Enzo Life Sciences), both reconstituted in 99.8% ethanol (Sigma-Aldrich), used at 10nM unless indicated. 99.8% ethanol was used as a carrier control at the same concentrations. Where indicated, IL-6 (Biolegend) or Tocilizumab (a kind gift from Dr Ceri Roberts in Professor Leonie Taams lab) were used in functional experiments.

### Flow cytometry and cell sorting

Cells were stained for 30 min at 4C, in 5mL polystyrene round bottom tubes (Falcon) in a final volume of 100μL staining buffer. For surface staining we used the following antibodies: CD4 (OKT4, Thermo Fisher Scientific), IL-6 (MQ2-13A5, Biologend), LAG-3 (3DS223H, Thermo Fisher Scientific), CD49b (P1H5, Thermo Fisher Scientific). DAPI or LIVE/DEAD Fixable Aqua/Violet (Invitrogen) staining were added to remove dead cells from the analysis. For intracellular staining cells, were activated with Phorbol 12-myristate 13-acetate (PMA) (50ng/mL, Sigma-Aldrich), ionomycin (1μg/mL, Sigma-Aldrich), GolgiStop (1X, BD Biosciences) under the presence of Brefeldin A (1X, BD Biosciences) for 5 hrs in fresh culture media at 37°C 5% CO2. Cells were then washed in PBS, initially stained for surface molecules and then fixed and permeabilized using Cytofix/Cytoperm buffer (eBioscience), followed by a wash in Perm/Wash buffer (eBioscience) and incubation with intracellular antibodies for 30 min at 4C. All samples were acquired on LSRFortessa (BD Biosciences) or Invitrogen Attune NxT Flow Cytometer within 24 hrs. Data was analyzed using FlowJo (LLC, v10.2). Assessment of cell proliferation by flow cytometry was carried out by CFSE dilution incorporation in polyclonally activated CD4^+^CD25^-^ T cells in presence of carrier or VitD after 72 hrs.

For FACS sorting of memory CD4^+^ T cells, bulk CD4^+^ T cells enriched as described using RosetteSep Human CD4^+^ enrichment cocktail were stained with antibodies against CD4 (OKT4, Thermo Fisher Scientific), CD45RA (HI100, Biolegend), CD45RO (UCLH1, BD Biosciences) and CD25 (2A3, BD Biosciences) in MACS buffer at 4C for 30 min. Following a wash in MACS buffer (0.5% bovine serum albumin (BSA;Sigma-Aldrich), 2mM EDTA (Sigma-Aldrich) in phosphate buffered saline (PBS;Gibco)), cells were re-suspended at a concentration of 30×10^6^ cells per mL and FACS sorted into CD4^+^CD25^-^ CD45RO^+^CD45RA^-^ memory cells to a purity of >99% using a BD FACSAria directly into culture medium.

### Cytokine measurement

Supernatants from 96-well plates were aliquoted and stored at −20C, after brief centrifugation to pellet cells. Cytokines were quantified using the Cytometric Bead Array (CBA) human/mouse Th1/Th2/Th17 kit (BD Biosciences) following manufacturers protocol using a FACSCanto II (BD Biosciences) and analyzed using FCAP Array software v3.0 (BD Biosciences). Alternatively, we used LEGENDplex Human Inflammation Panel 1 (13plex) (740808, BioLegend) following manufacturers protocol using an Invitrogen Attune NxT Flow Cytometer and analyzed using FlowJo v9 software (BD Biosciences).

### Western blot

Whole cell extracts were prepared by cell lysis in RIPA buffer (Thermo Fisher Scientific), as per manufacturer’s protocol, with added Protease Inhibitor Cocktail Set III (Calbiochem), at a 1 in 250 dilution, and 5% 2-Mercaptoethanol, then denatured at 95C for 5 min. Protein concentration was quantified using a quickstart protein assay (BioRad). Proteins were resolved by SDS-PAGE on 10% Tris-Glycine gels (Invitrogen) and electro transferred onto polyvinylidene fluoride membranes (Immobilon, Millipore). Immunoblotting was performed according to standard protocols with initial blocking in PBS with 10% w/v clotting grade blocker (BioRad) or 3% w/v BSA (Sigma-Aldrich) and 0.1% v/v Tween20 (Sigma-Aldrich), followed by overnight incubation in primary antibodies: anti-VDR (D-6, Santa Cruz biotechnology) pSTAT3 (D3A7, Cell Signaling Technologies), STAT3 (124H6, Cell Signaling Technologies), Phospho-c-Jun (S63, R&D systems) or c-Jun (L70B11, Cell Signaling Technologies), followed by blocking and two hours incubation in appropriate HRP conjugated secondary anti-mouse or anti-rabbit antibodies (TrueBlot antibodies, Rockland). Bound antibodies were detected using ECL-Plus reagents (Thermo Fisher Scientific) and visualized on an imageQUant LAS4000 mini (GE Healthcare). Blots were quantified using ImageStudioLite (LICOR).

Nuclear and cytoplasmic extracts were separated using NE-PER™ Nuclear and Cytoplasmic Extraction (Thermo Fisher Scientific) reagents as per manufacturer’s instructions. Briefly, 1×10^6^ CD4^+^CD25^-^ cells following 48 hr cell culture, as indicated, were pelleted and lysed in CER II to obtain cytoplasmic extract and NER to obtain nuclear extract. Western botting was then carrier out as specified above using anti-Hsp90 (C45G5, Cell Signaling Technologies) and anti-histone H2A.X (D17A3, Cell Signaling Technologies).

For BACH2 Western blotting, snap-frozen cell pellets of 600,000 cells were resuspended directly in 2X Laemmli Sample Buffer (Bio-Rad) supplemented with 5% 2-Mercaptoethanol, denatured at 95C for 5 min, and separated using a kD Criterion TGX gel (Bio-Rad) at 120V. Proteins were then transferred onto nitrocellulose using a Trans-Blot TurboTM Midi Nitrocellulose Transfer Pack with the Trans-Blot TurboTM Transfer System as per manufacturer’s guidelines (Bio-Rad). Membranes were incubated in Odyssey Blocking Buffer with PBS (LI-COR Biosciences) for 1 hr at RT prior to the addition of primary antibodies. Antibody against BACH2 (D3T3G, Cell Signaling Technologies, 1:5000 dilution) was added in blocking buffer and incubated with membranes overnight at 4C with rotation. Membranes were then washed thrice in wash buffer (PBS with 0.1% Tween-20) and exposed to secondary antibody (LI-COR Biosciences, IRDye 800CW, goat anti-rabbit green 926-32211) at a dilution of 1:5000 in blocking buffer for 1 hr at RT. After three washes, protein signals were visualized using the Odyssey CLx (LI-COR Biosciences). The membrane was re-probed with anti-β-Tubulin (2146, Cell Signaling Technologies, 1:1000 dilution). Quantifications were performed in Image Studio v5.2 (LI-COR Biosciences).

### Human phospho-kinase antibody array

Array (Proteome Profiler Human Phospho-Kinase Array Kit, R&D systems) was carried out as per manufacturer’s protocol. Briefly, CD4+CD25- T cells from two donors were activated in the presence of ethanol carrier or VitD, lysed in manufacturer’s lysis buffer and protein concentrations were quantified using quickstartTM protein assay (BioRad) and concentrations adjusted to 800μg/mL before 334μL were loaded on each membrane. Signals were amplified by Chemi-Reagent Mix and visualized using an imageQuant LAS4000 mini (GE Healthcare) and quantified by ImageStudioLite (LICOR). Reference spots on each membrane were used to normalize signals across membranes.

### Quantification of VDR and DAPI colocalization

VDR and DAPI colocalization was performed in memory CD4^+^ T cells stained with mouse anti-human VDR (D-6, Santa Cruz), anti-mouse-AlexaFluor647 and DAPI. Data was acquired on an ImageStreamX system equipped with three lasers, running Inspire software and analyzed with IDEAS 3.0 software (all from AMNIS Corp, Seattle, WA, USA). Co-localization between VDR and DAPI was assessed with specific features/masks within IDEAS 3.0. A minimum of 100,000 cells/events were acquired per sample.

### Quantitative PCR

Cells (typically 2-4×10^6^) were lysed in 350μl Trizol reagent (Ambion, Life Technologies) and RNA extracted using the Direct-zol RNA Miniprep kit (Zymo) with on column genomic DNA digestion, according to manufacturer’s instructions. RNA concentration was assessed using a NanoDrop (NanoDrop) spectrophotometer. RNA was then reverse transcribed to cDNA using the qPCRBIO cDNA Synthesis Kit (PCR Biosystems) according to the manufacturer’s suggestion. qPCR was carried out in a 384-well plate using the ViiA7 real-time PCR system (Life technologies) and SYBR Green PCR Master Mix (Life Technologies). All reactions were carried out in triplicate using 18s and UBC (geNorm 6 gene kit, catalogue number ge-SY-6 from PrimerDesign UK) as housekeeping genes. Gene specific primers for IL-6 (Hs_IL6_1_SG QuantiTect Primer Assay) was sourced from QuantiTect (Qiagen) primers. qPCR for *CYP27B1* (Hs01096154_m1) and *VDR* (Hs01045843_m1), with 18S (Hs99999901_s1) as housekeeping control were carried out on RNA reverse transcribed to cDNA with an iScript cDNA synthesis kit (Bio-Rad), using Taqman chemistry and gene-specific probes purchased from Applied Biosystems. Melt curve analysis to ensure amplification of a single product was used. Comparative Ct method was used for analysis using the Viia7 software producing a Ct (relative quantity (RQ) compared to a reference sample).

### STAT3 inhibition

For STAT3 knock-down, memory CD4^+^ T cells were sorted and rested overnight in culture medium at 37C 5%CO2. The next day, 5 million cells were washed twice with PBS and the cell pellet was nucleofected with 10nM Silencer Select STAT3 s745 (sequence for sense GCACCUUCCUGCUAAGAUUTT and antisense AAUCUUAGCAGGAAGGUGCCT) or Silencer Select Negative Control (sequence proprietary) siRNA (Thermo Fisher Scientific) using the Amaxa Human T Cell Nucleofector Kit (Lonza) and the Nucleofector 2b Device (Lonza) (U-014 program). After nucleofection, cells were rested in culture media at 37C 5%CO2 for 6 hrs. Dead cells were removed with Ficoll-Hypaque density gradient centrifugation (Lymphoprep, Axis-Shield, Oslo, Norway) and 2×10^6^ cells were cultured with anti-CD3/CD28 beads (Thermo Fisher Scientific) at a cell:bead ratio of 4:1, with VitD or carrier, for 72 hrs at 37C 5%CO2.

For chemical STAT3 inhibition, memory CD4^+^ T cells were sorted and rested overnight in culture medium at 37C 5%CO2. Next day, 2×10^6^ cells were activated anti-CD3/CD28 beads (Thermo Fisher Scientific) in a 4:1 ratio of cells:beads for 72 hrs at 37°C, 5%CO2 with VitD or carrier in the presence of 150 nM Curcubitacin I, *Cucumis sativus L*. (Calbiochem) or carrier (ethanol).

### H3K27Ac and c-JUN CUT&RUN

*In situ* CUT&RUN sequencing was performed on memory CD4^+^ T cells activated with anti-CD3+anti-CD28 in the presence of carrier or VitD for 0.75 hr, 48 hrs and 72 hrs using the published protocol of Skene et al. ^48^. Rabbit antibodies targeting H3K27Ac (ab4729, Abcam), c-JUN (60AB, Cell Signaling Technologies), and non-specific IgG (31235, Thermo Fisher Scientific) were used, together with pAG-MNase (123461, Addgene) for small DNA fragment generation. All chemicals required for CUT&RUN buffers were purchased from Sigma-Aldrich and Thermo Fisher Scientific. Once in solution, isolated DNA fragments were then immediately prepared for Illumina sequencing (NEB Ultra II, New England Biolabs) as per the manufacturer’s suggestions. Amplified libraries were quantified by high sensitivity fluorometry (DeNovix) and high sensitivity D1000 tape analysis using a 4200 TapeStation (Agilent Technologies).

For H3K27Ac samples, the alignment of sequenced reads to the human reference genome (GRCh37; hg19) was conducted using Bowtie2^68^ with default pair-end alignment settings and additional parameters ‘--local --maxins 250’. Sorting and indexing of the aligned reads were conducted by samtools/1.10^69^. Reads with a mapping quality less than 30 were discarded. Binding peaks were called by ‘callpeaks’ procedure from MACS2^70^ using default parameters except ‘-f BAMPE --nomodel’. For JUN samples, all sequencing reads were mapped to the human reference genome (hg19) without ‘--local’ parameter (end-to-end). Sequencing fragments with length less than 120 bp were further sorted and indexed using samtools/1.10^69^. For discrimination of JUN induced peaks, MACS2 parameters ‘--nomodel -g hs -f BAMPE -q 0.01 --SPMR --keep-dup all’ was applied. IgG BAM files were included as controls for MACS2 peak calling. The identified peaks for H3K27Ac and JUN experiments were further screened against ‘hg19 blacklisted’ genomic regions, mitochondrial DNA, and pseudo-chromosomes. H3K27ac and JUN mean binding intensities in peak regions were calculated by UCSC bigWigAverageOverBed with default parameters. Peaks, combined from two conditions, with mean binding intensity greater than 0.2 in at least one condition and fold change of intensity greater than 1.5 were defined as differential peaks. Known motif finding on identified differential peaks (+/-100bp around the peak summit) was done by Homer findMotifsGenome.pl using parameter ‘-size given’. *De novo* motif footprinting for c-JUN CUT&RUN was performed with CutRunTools^71^ and the resulting cut matrices were used to compute average cuts per bp in proximity to the de novo called JUN motif. CUT&RUN tracks and heatmaps were visualized using IGV browser (Broad Institute) and deepTools^72^, respectively.

### *De novo* super-enhancer calling

Super-Enhancers were called using ROSE (Rank Ordering of Super-Enhancers) under default settings^49^. Briefly, H3K27Ac CUT&RUN VitD induced peaks were selected (see above), BAM files were sorted and indexed using samtools/1.10^69^, then sorted VitD and carrier H3K27Ac BAM files were used as additional input for ROSE_main.py. Carrier was denoted as control. SE annotations were manually screened and corrected, if needed. Hockey-stick plots of rank ordered stitched-enhancers plotted against VitD minus carrier H3K27Ac signal were produced in RStudio (ver. 1.2.5033)

### RNA sequencing and transcriptome analysis

Memory CD4^+^ T cells, including from BACH2^WT/L24P^ patient samples, were isolated and activated with or without VitD treatment as described. 6×10^5^ unactivated cells, or cells activated for 48 hrs, were pelleted at 300g for 5 min. RNA was stabilized by lysing cells in 100μL of RNAqueous lysis buffer and snap-freezing on dry-ice followed by storage at −80C followed by isolation of total RNA by the RNAqueous^®^-Micro kit (AM1931, Thermo Fisher Scientific) as per manufacturers guidelines. 1μg total RNA for each sample was subjected to NEBNext Poly(A) mRNA Magnetic Isolation Module (E7490). Resulting mRNA was prepared for RNA-seq using a NEB Ultra II RNA Sequencing Library Prep Kit (New England Biolabs) as per the manufacturer’s guidelines.

The expression level of all genes in all RNA-seq libraries was quantified by ‘rsem-calculate-expression’ in RSEM^73^ v1.3.1 with parameters ‘--bowtie-n 1 --bowtie-m 100 --seed-length 28 --bowtie-chunkmbs 1000’. The bowtie index for RSEM alignment was generated by ‘rsem-prepare-reference’ on all RefSeq genes, downloaded from UCSC table browser in April 2017. EdgeR^74^ v3.26.8 was used to normalize gene expression among all libraries and identify differentially expressed genes among samples. Analysis of microarray data sourced from GSE119416 was carried out using Partek^®^ Genomics Suite^®^ (Partek, Inc.). Differentially expressed gene thresholds were defined using the following criteria: at least 1.5-fold change in either direction at p-value<0.05 for microarray analysis; at least 1.75-fold change in either direction at FDR<0.05 for RNA-seq analysis.

### Single cell RNA-sequencing studies and analysis

#### COVID-19 bronchoalveolar lavage dataset

The pre-processed h5 matrix files for nine COVID-19 patient bronchoalveolar-lavage (BAL) samples and three healthy control BAL samples were obtained from GSE145926^75^ and GSE122960^76^, respectively. Read mapping and basic filtering were performed with the Cell Ranger pipeline by the original authors. We further processed the samples using Seurat (version 3) as follows: Only genes found to be expressed in more than 3 cells were retained. Cells with gene number between 200 and 6,000, UMI count >1,000 were retained. Cells with >10% of their unique molecular identifiers (UMIs) mapping to mitochondrial genes or cells with <300 features were discarded to eliminate low quality cells. This yielded a total of 66453 cells across 12 samples. The filtered count matrices were then normalized by total UMI counts, multiplied by 10,000 and transformed to natural log space. The top 2000 variable features were determined based on the variance stabilizing transformation function (FindVariableFeatures) by Seurat using default parameters. All samples were integrated using canonical correlation analysis (FindIntegrationAnchors/IntegrateData) function with parameter k.filter=140. Variants arising from library size and percentage of mitochondrial genes were regressed out by the ScaleData function in Seurat. Principal Component Analysis (PCA) was performed and the top 50 Principal Components (PCs) were included in a Uniform Manifold Approximation and Projection (UMAP) dimensionality reduction. Clusters were identified on a shared nearest neighbor (SNN) modularity graph using the top 50 PCs and the original Louvain algorithm. Cluster annotations were based on canonical marker genes.

Cells identified as “T” and “Cytotoxic T” were subsetted and reprocessed in Seurat. Samples C146 and C52 were removed due to low number of T cells (19 and 56 cells, respectively). Integration was performed using FindIntegrationAnchors/IntegrateData with following parameters: “dims=1:30 k.filter=124”. Normalization, variable feature detection, scaling, dimensionality reduction and clustering were performed, similar to that described above, using the top 30 PCs and clustering resolution of 0.3. Annotation was guided by canonical marker genes (**Fig. S1c**). The contaminating macrophage cluster annotated by canonical macrophage marker genes (e.g. CD14) was removed from the analysis. Statistical differences were assessed by two-tailed Wilcoxon test. Differentially expressed genes were defined by fold change of >1.5, adjusted p value of <0.05 and expression in at least 10% of cells in either comparison cluster.

Downstream analysis was focused on cells annotated as CD4^+^ T cells (cluster CD4^+^ T (A) and CD4^+^ T (B) combined). DEGs between COVID-19 and Healthy CD4^+^ T cells were visualized using Morpheus (https://software.broadinstitute.org/morpheus/). Genes upregulated in COVID-19 CD4^+^ T cells were subjected to pathway analysis (**Fig. 1c**) using hallmark geneset collection from Molecular Signatures Database v7.1 (MSigDB) or drug prediction from National Toxicology Program’s DrugMatrix genesets (n=7876) using EnrichR R package (**Fig. 4f**). Gene list scores were calculated by the AddModuleScore function in Seurat^77^ with a control gene set size of 100. The correlation and the ROC analyses on the module scores comparing patients and healthy controls were done in R or Prism 8.4.0.

#### COVID-19 PBMC dataset

The pre-processed serialized R objects for six COVID-19 patient PBMC samples and six healthy control PBMC samples were obtained from GSE150728^78^. Read mapping and basic filtering were performed with the Cell Ranger pipeline (10x genomics) by the original authors. The exonic count matrices were further processed by Seurat (version 3) as follows: Only genes found to be expressed in more than 10 cells were retained. The QC steps for filtering the samples were performed as described^78^. Briefly, cells with 1,000-15,000 UMIs and <20% of reads from mitochondrial genes were retained. Cells with >20% of reads mapped to RNA18S5 or RNA28S5, and/or expressed more than 75 genes per 100 UMIs were excluded. SCTransform function was invoked to normalize the dataset and to identify variable genes as previously described^78^. PCA was performed and the top 50 PCs were included in a UMAP dimensionality reduction. Clusters were identified on a SNN modularity graph using the top 50 PCs and the original Louvain algorithm. Cluster annotations were based on canonical marker genes. Gene list module scores were calculated by the AddModuleScore function in Seurat^77^ with a control gene set size of 100.

### Geneset Enrichment Analysis (GSEA) and gene lists used in the manuscript

GSEA was carried out using published methods^55^. Tr1 gene lists were obtained from GSE139990. Data were filtered to include only genes expressed at greater than 0.25 counts per million in at least 2 samples, TMM normalized within the edgeR package (v3.28.1), and differential expression performed using the glmQLFit and glmQLFTest functions in edgeR. Tr1 signatures were then defined as genes differentially expressed (at least 4-fold change in either direction at FDR<0.05) between Tr1 and Th0 cells. Th1/Th2/Th17 gene lists were obtained from transcriptomes of sorted mouse CD4^+^ T cell subsets^79^. VitD-regulated genes were defined as described in RNA-sequencing methods. BACH2 bound and regulated genes were obtained from GSE45975^6^. All gene lists used in this manuscript are shown in **Table S7**.

### Psoriasis Lesional Skin Staining, Confocal Imaging, and Analysis

All study participants were provided written informed consent and data for all psoriasis patients were obtained under the Psoriasis, Atherosclerosis and Cardiometabolic Disease Initiative (PACI, 13-H-0065). The sample population included 3 psoriasis patients not on VitD supplementation and 2 psoriasis patients on VitD supplementation. 3 mm punch-biopsies were collected from psoriasis lesional skin and placed directly into 10% formalin. Lesional skin samples were sectioned into 7μm sections and mounted onto glass slides. The samples were de-waxed, and antigens were retrieved using the citrate buffer method. Samples were washed once in PBS and blocked for 20 min at RT in 10% normal goat serum. The samples were incubated with anti-CD3 (F7.2.38, Thermo Fisher Scientific) and anti-BACH2 (D3TG, Cell Signaling) primary antibodies overnight at 4C and washed in 0.01M PBS the next day. Subsequently, Alexa Fluor 488 (Jackson Immuno Research Laboratories Inc., 115-546-062) and Alexa Fluor 594 (Jackson Immuno Research Laboratories Inc., 111-586-047) secondary antibodies were added for 1 hr at RT and then washed in 0.01M PBS. Hoechst was added to the samples for 10 min, samples were washed and then cover slipped with Fluoromount G. Confocal images were collected on a Zeiss 780 inverted confocal microscope at 40X with oil-immersion and analyzed using ImageJ software. Representative confocal images were prepared in Zen Blue 3.1 (Zeiss). For each sample, 5-6 images were acquired. In order to quantify the frequency of immune cells in the dermis, images were cropped to 1200 x 1200 pixels to exclude the epidermis and the number of nuclei in the 408 nm channel were determined by manual thresholding in ImageJ. Subsequent fluorescent analysis was completed to determine the number of nuclei per image. The number of BACH2 positive cells in the 594 nm channel were calculated per image and the number of BACH2 foci per BACH2 positive cells were recorded.

### Data presentation and statistical analysis

Figures were prepared using Adobe Illustrator (Adobe). Statistical analysis and graphical visualizations were carried out in GraphPad Prism (v.8.4.0), XLstat biomed (v2017.4), DataGraph 4.5.1 (Visual Data Tools, Inc.), Cytoscape 3^80^ and Circos Table Viewer v.90.63.9^81^. Statistical analyses were carried out using appropriate paired or unpaired parametric and non-parametric tests, as required. Multiple comparisons were performed using ANOVA. p-values of <0.05 were considered statistically significant throughout.

### Data availability statement

The data generated for this study will be deposited at the Gene Expression Omnibus (GEO) and freely available.

## Notes

### Competing Interest Statement

The authors have declared no competing interest.

## References

1. Neyer, L. E. et al. Role of interleukin-10 in regulation of T-cell-dependent and T-cell-independent mechanisms of resistance to Toxoplasma gondii. Infect Immun 65, 1675–1682 (1997).

2. Hunter, C. A. et al. IL-10 is required to prevent immune hyperactivity during infection with Trypanosoma cruzi. J Immunol 158, 3311–3316 (1997).

3. West, E. E., Kolev, M. & Kemper, C. Complement and the Regulation of T Cell Responses. Annu Rev Immunol 36, 309–338 (2018).

4. Yan, B. et al. SARS-CoV2 drives JAK1/2-dependent local and systemic complement hyper-activation. ResearchSquare (2020). doi:10.21203/rs.3.rs-33390/v1

5. Vahedi, G. et al. Super-enhancers delineate disease-associated regulatory nodes in T cells. Nature 520, 558–562 (2015).

6. Roychoudhuri, R. et al. BACH2 represses effector programs to stabilize T(reg)-mediated immune homeostasis. Nature 498, 506–510 (2013).

7. Afzali, B. et al. BACH2 immunodeficiency illustrates an association between super-enhancers and haploinsufficiency. Nat Immunol 18, 813–823 (2017).

8. Zhou, F. et al. Clinical course and risk factors for mortality of adult inpatients with COVID-19 in Wuhan, China: a retrospective cohort study. Lancet 395, 1054–1062 (2020).

9. Horby, P. et al. Effect of Dexamethasone in Hospitalized Patients with COVID-19: Preliminary Report. medRxiv 2020.06.22.20137273 (2020). doi:10.1101/2020.06.22.20137273

10. Spagnolo, P. et al. Pulmonary fibrosis secondary to COVID-19: a call to arms? Lancet Respir Med (2020). doi:10.1016/S2213-2600(20)30222-8

11. Carfì, A., Bernabei, R., Landi, F. Gemelli Against COVID-19 Post-Acute Care Study Group. Persistent Symptoms in Patients After Acute COVID-19. JAMA (2020). doi:10.1001/jama.2020.12603

12. Zhao, J. et al. Airway Memory CD4(+) T Cells Mediate Protective Immunity against Emerging Respiratory Coronaviruses. Immunity 44, 1379–1391 (2016).

13. Grifoni, A. et al. Targets of T Cell Responses to SARS-CoV-2 Coronavirus in Humans with COVID-19 Disease and Unexposed Individuals. Cell 181, 1489–1501.e15 (2020).

14. Lucas, C. et al. Longitudinal immunological analyses reveal inflammatory misfiring in severe COVID-19 patients. medRxiv 2020.06.23.20138289 (2020). doi:10.1101/2020.06.23.20138289

15. Kolev, M. et al. Diapedesis-Induced Integrin Signaling via LFA-1 Facilitates Tissue Immunity by Inducing Intrinsic Complement C3 Expression in Immune Cells. Immunity 52, 513–527.e8 (2020).

16. Magro, C. et al. Complement associated microvascular injury and thrombosis in the pathogenesis of severe COVID-19 infection: a report of five cases. Transl Res (2020). doi:10.1016/j.trsl.2020.04.007

17. Cardone, J. et al. Complement regulator CD46 temporally regulates cytokine production by conventional and unconventional T cells. Nat Immunol 11, 862–871 (2010).

18. Cope, A., Le Friec, G., Cardone, J. & Kemper, C. The Th1 life cycle: molecular control of IFN-γ to IL-10 switching. Trends Immunol 32, 278–286 (2011).

19. Meyer, M. B. & Pike, J. W. Mechanistic homeostasis of vitamin D metabolism in the kidney through reciprocal modulation of Cyp27b1 and Cyp24a1 expression. The Journal of Steroid Biochemistry and Molecular Biology 196, 105500 (2020).

20. Sigmundsdottir, H. et al. DCs metabolize sunlight-induced vitamin D3 to ‘program’ T cell attraction to the epidermal chemokine CCL27. Nat Immunol 8, 285–293 (2007).

21. Veldman, C. M., Cantorna, M. T. & DeLuca, H. F. Expression of 1,25-dihydroxyvitamin D(3) receptor in the immune system. Arch. Biochem. Biophys. 374, 334–338 (2000).

22. Liszewski, M. K. et al. Intracellular complement activation sustains T cell homeostasis and mediates effector differentiation. Immunity 39, 1143–1157 (2013).

23. van Etten, E. & Mathieu, C. Immunoregulation by 1,25-dihydroxyvitamin D3: basic concepts. The Journal of Steroid Biochemistry and Molecular Biology 97, 93–101 (2005).

24. Martineau, A. R. et al. Vitamin D supplementation to prevent acute respiratory tract infections: systematic review and meta-analysis of individual participant data. BMJ 356, i6583 (2017).

25. Yamshchikov, A. V., Desai, N. S., Blumberg, H. M., Ziegler, T. R. & Tangpricha, V. Vitamin D for treatment and prevention of infectious diseases: a systematic review of randomized controlled trials. Endocr Pract 15, 438–449 (2009).

26. Holick, M. F. Vitamin D deficiency. N Engl J Med 357, 266–281 (2007).

27. Chen, D.-J. et al. Altered microRNAs expression in T cells of patients with SLE involved in the lack of vitamin D. Oncotarget 8, 62099–62110 (2017).

28. Jeffery, L. E. et al. 1,25-Dihydroxyvitamin D3 and IL-2 combine to inhibit T cell production of inflammatory cytokines and promote development of regulatory T cells expressing CTLA-4 and FoxP3. J Immunol 183, 5458–5467 (2009).

29. Villegas-Ospina, S., Aguilar-Jimenez, W., Gonzalez, S. M. & Rugeles, M. T. Vitamin D modulates the expression of HLA-DR and CD38 after in vitro activation of T-cells. Horm Mol Biol Clin Investig 29, 93–103 (2017).

30. Berge, T. et al. The multiple sclerosis susceptibility genes TAGAP and IL2RA are regulated by vitamin D in CD4+ T cells. Genes Immun 17, 118–127 (2016).

31. Groux, H. et al. A CD4+ T-cell subset inhibits antigen-specific T-cell responses and prevents colitis. Nature 389, 737–742 (1997).

32. Barrat, F. J. et al. In vitro generation of interleukin 10-producing regulatory CD4(+) T cells is induced by immunosuppressive drugs and inhibited by T helper type 1 (Th1)- and Th2-inducing cytokines. J Exp Med 195, 603–616 (2002).

33. Gagliani, N. et al. Coexpression of CD49b and LAG-3 identifies human and mouse T regulatory type 1 cells. Nat Med 19, 739–746 (2013).

34. Hunter, C. A. & Jones, S. A. IL-6 as a keystone cytokine in health and disease. Nat Immunol 16, 448–457 (2015).

35. Stumhofer, J. S. et al. Interleukins 27 and 6 induce STAT3-mediated T cell production of interleukin 10. Nat Immunol 8, 1363–1371 (2007).

36. Jin, J.-O., Han, X. & Yu, Q. Interleukin-6 induces the generation of IL-10-producing Tr1 cells and suppresses autoimmune tissue inflammation. J Autoimmun 40, 28–44 (2013).

37. Bettelli, E. et al. Reciprocal developmental pathways for the generation of pathogenic effector TH17 and regulatory T cells. Nature 441, 235–238 (2006).

38. Mangan, P. R. et al. Transforming growth factor-beta induces development of the T(H)17 lineage. Nature 441, 231–234 (2006).

39. Yegorov, S., Bromage, S., Boldbaatar, N. & Ganmaa, D. Effects of Vitamin D Supplementation and Seasonality on Circulating Cytokines in Adolescents: Analysis of Data From a Feasibility Trial in Mongolia. Front Nutr 6, 166 (2019).

40. Turksen, K., Kupper, T., Degenstein, L., Williams, I. & Fuchs, E. Interleukin 6: insights to its function in skin by overexpression in transgenic mice. Proc Natl Acad Sci USA 89, 5068–5072 (1992).

41. Lin, Z.-Q., Kondo, T., Ishida, Y., Takayasu, T. & Mukaida, N. Essential involvement of IL-6 in the skin wound-healing process as evidenced by delayed wound healing in IL-6-deficient mice. J. Leukoc. Biol. 73, 713–721 (2003).

42. Laurent, S., Le Parc, J.-M., Clérici, T., Bréban, M. & Mahé, E. Onset of psoriasis following treatment with tocilizumab. Br. J. Dermatol. 163, 1364–1365 (2010).

43. Wendling, D., Letho-Gyselinck, H., Guillot, X. & Prati, C. Psoriasis onset with tocilizumab treatment for rheumatoid arthritis. The Journal of Rheumatology 39, 657 (2012).

44. Herdick, M. & Carlberg, C. Agonist-triggered modulation of the activated and silent state of the vitamin D(3) receptor by interaction with co-repressors and co-activators. J. Mol. Biol. 304, 793–801 (2000).

45. Oda, Y., Chalkley, R. J., Burlingame, A. L. & Bikle, D. D. The transcriptional coactivator DRIP/mediator complex is involved in vitamin D receptor function and regulates keratinocyte proliferation and differentiation. J Invest Dermatol 130, 2377–2388 (2010).

46. Polly, P. et al. VDR-Alien: a novel, DNA-selective vitamin D(3) receptor-corepressor partnership. FASEB J 14, 1455–1463 (2000).

47. Wei, Z. et al. Vitamin D Switches BAF Complexes to Protect β Cells. Cell 173, 1135–1149.e15 (2018).

48. Skene, P. J., Henikoff, J. G. & Henikoff, S. Targeted in situ genome-wide profiling with high efficiency for low cell numbers. Nature Protocols 13, 1006–1019 (2018).

49. Whyte, W. A. et al. Master Transcription Factors and Mediator Establish Super-Enhancers at Key Cell Identity Genes. Cell 153, 307–319 (2013).

50. Lovén, J. et al. Selective inhibition of tumor oncogenes by disruption of super-enhancers. Cell 153, 320–334 (2013).

51. Bohmann, D. et al. Human proto-oncogene c-jun encodes a DNA binding protein with structural and functional properties of transcription factor AP-1. Science 238, 1386–1392 (1987).

52. Povoleri, G. A. M. et al. Human retinoic acid-regulated CD161+ regulatory T cells support wound repair in intestinal mucosa. Nat Immunol 19, 1403–1414 (2018).

53. Cooper, J. D. et al. Meta-analysis of genome-wide association study data identifies additional type 1 diabetes risk loci. Nat Genet 40, 1399–1401 (2008).

54. Franke, A. et al. Genome-wide meta-analysis increases to 71 the number of confirmed Crohn’s disease susceptibility loci. Nat Genet 42, 1118–1125 (2010).

55. Subramanian, A. et al. Gene set enrichment analysis: a knowledge-based approach for interpreting genome-wide expression profiles. Proc Natl Acad Sci USA 102, 15545–15550 (2005).

56. Armstrong, A. W. & Read, C. Pathophysiology, Clinical Presentation, and Treatment of Psoriasis: A Review. JAMA 323, 1945–1960 (2020).

57. Tashiro, S. et al. Repression of PML nuclear body-associated transcription by oxidative stress-activated Bach2. Mol Cell Biol 24, 3473–3484 (2004).

58. Laird, E., Rhodes, J. & Kenny, R. A. Vitamin D and Inflammation: Potential Implications for Severity of Covid-19. Ir Med J 113, 81 (2020).

59. Ilie, P. C., Stefanescu, S. & Smith, L. The role of vitamin D in the prevention of coronavirus disease 2019 infection and mortality. Aging Clin Exp Res (2020). doi:10.1007/s40520-020-01570-8

60. Davies, G., Garami, A. R. & Byers, J. C. Evidence Supports a Causal Model for Vitamin D in COVID-19 Outcomes. medRxiv 2020.05.01.20087965 (2020). doi:10.1101/2020.05.01.20087965

61. De Smet, D., De Smet, K., Herroelen, P., Gryspeerdt, S. & Martens, G. A. Vitamin D deficiency as risk factor for severe COVID-19: a convergence of two pandemics. medRxiv 2020.05.01.20079376 (2020). doi:10.1101/2020.05.01.20079376

62. Daneshkhah, A. et al. The Possible Role of Vitamin D in Suppressing Cytokine Storm and Associated Mortality in COVID-19 Patients. medRxiv 2020.04.08.20058578 (2020). doi: 10.1101/2020.04.08.20058578

63. Moozhipurath, R. K., Kraft, L. & Skiera, B. Evidence of Protective Role of Ultraviolet-B (UVB) Radiation in Reducing COVID-19 Deaths. medRxiv 2020.05.06.20093419 (2020). doi:10.1101/2020.05.06.20093419

64. Liberzon, A. et al. Molecular signatures database (MSigDB) 3.0. Bioinformatics 27, 1739–1740 (2011).

65. Liberzon, A. et al. The Molecular Signatures Database (MSigDB) hallmark gene set collection. Cell Syst 1, 417–425 (2015).

66. Hidalgo, A. A., Deeb, K. K., Pike, J. W., Johnson, C. S. & Trump, D. L. Dexamethasone enhances 1alpha, 25-dihydroxyvitamin D3 effects by increasing vitamin D receptor transcription. J Biol Chem 286, 36228–36237 (2011).

## Methods only references

67. Kolev, M. et al. Complement Regulates Nutrient Influx and Metabolic Reprogramming during Th1 Cell Responses. Immunity 42, 1033–1047 (2015).

68. Langmead, B. & Salzberg, S. L. Fast gapped-read alignment with Bowtie 2. Nature Methods 9, 357–359 (2012).

69. Li, H. et al. The Sequence Alignment/Map format and SAMtools. Bioinformatics 25, 2078–2079 (2009).

70. Zhang, Y. et al. Model-based analysis of ChIP-Seq (MACS). Genome Biol 9, R137 (2008).

71. Zhu, Q., Liu, N., Orkin, S. H. & Yuan, G.-C. CUT&RUNTools: a flexible pipeline for CUT&RUN processing and footprint analysis. Genome Biol 20, 192 (2019).

72. Ramírez, F. et al. deepTools2: a next generation web server for deep-sequencing data analysis. Nucleic Acids Res 44, W160–5 (2016).

73. Li, B. & Dewey, C. N. RSEM: accurate transcript quantification from RNA-Seq data with or without a reference genome. BMC Bioinformatics 12, 323 (2011).

74. Robinson, M. D., McCarthy, D. J. & Smyth, G. K. edgeR: a Bioconductor package for differential expression analysis of digital gene expression data. Bioinformatics 26, 139–140 (2010).

75. Liao, M. et al. Single-cell landscape of bronchoalveolar immune cells in patients with COVID-19. Nat Med 19, 181 (2020).

76. Reyfman, P. A. et al. Single-Cell Transcriptomic Analysis of Human Lung Provides Insights into the Pathobiology of Pulmonary Fibrosis. Am. J. Respir. Crit. Care Med. 199, 1517–1536 (2019).

77. Tirosh, I. et al. Dissecting the multicellular ecosystem of metastatic melanoma by single-cell RNA-seq. Science 352, 189–196 (2016).

78. Wilk, A. J. et al. A single-cell atlas of the peripheral immune response in patients with severe COVID-19. Nat Med (2020). doi:10.1038/s41591-020-0944-y

79. Stubbington, M. J. et al. An atlas of mouse CD4(+) T cell transcriptomes. Biol. Direct 10, 14 (2015).

80. Shannon, P. et al. Cytoscape: a software environment for integrated models of biomolecular interaction networks. Genome Res. 13, 2498–2504 (2003).

81. Krzywinski, M. et al. Circos: an information aesthetic for comparative genomics. Genome Res. 19, 1639–1645 (2009).

