## Supplementary Figures for "An autocrine Vitamin D-driven Th1 shutdown program can be exploited for COVID-19"

<u>List of supplementary data</u>

Supplementary Figures 1-11
Supplementary Tables 1-7

### Fig. S1

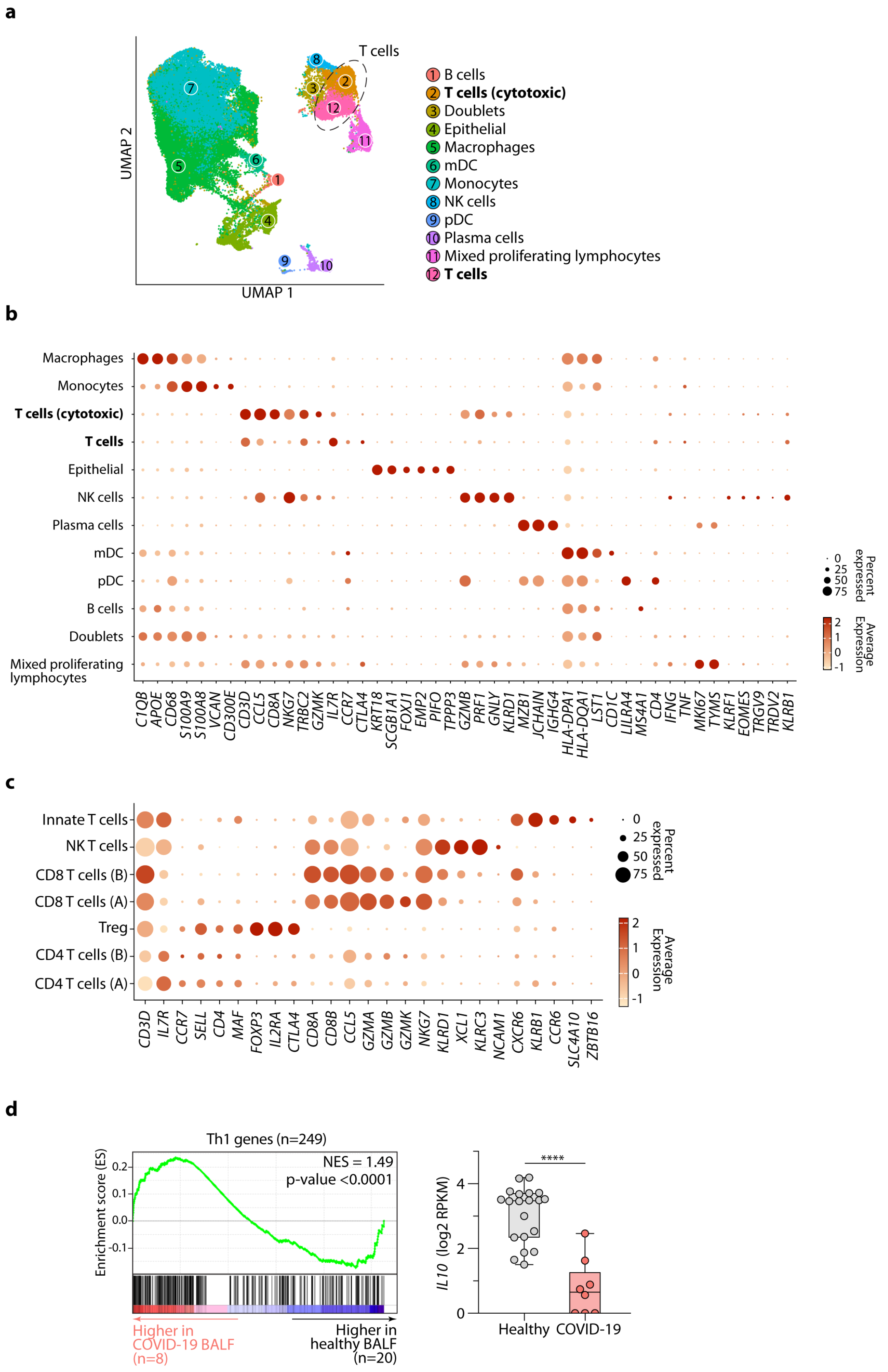

a
T cells
UMAP 2
UMAP 1
1 B cells
2 T cells (cytotoxic)
3 Doublets
4 Epithelial
5 Macrophages
6 mDC
7 Monocytes
8 NK cells
9 pDC
10 Plasma cells
11 Mixed proliferating lymphocytes
12 T cells
b
Macrophages
Monocytes
T cells (cytotoxic)
T cells
Epithelial
NK cells
Plasma cells
mDC
pDC
B cells
Doublets
Mixed proliferating lymphocytes
C1QB APOE CD68 S100A9 S100A8 VCAN CD300E CD3D CCL5 CD8A NKG7 TRBC2 GZMK IL7R CCR7 CTLA4 KRT18 SCGB1A1 FOXJ1 EMP2 PIFO TPPP3 GZMB PRF1 GNLY KLRD1 MZB1 JCHAIN IGHG4 HLA-DPA1 HLA-DQA1 LST1 CD1C LILRA4 MS4A1 CD4 IFNG TNF MKI67 TYMS KLRF1 EOMES TRGV9 TRDV2 KLRB1
Percent expressed
0
25
50
75
Average Expression
2
1
0
-1
c
Innate T cells
NK T cells
CD8 T cells (B)
CD8 T cells (A)
Treg
CD4 T cells (B)
CD4 T cells (A)
CD3D IL7R CCR7 SELL CD4 MAF FOXP3 IL2RA CTLA4 CD8A CD8B CCL5 GZMA GZMB GZMK NKG7 KLRD1 XCL1 KLRC3 NCAM1 CXCR6 KLRB1 CCR6 SLC4A10 ZBTB16
Percent expressed
0
25
50
75
Average Expression
2
1
0
-1
d
Th1 genes (n=249)
NES = 1.49
p-value <0.0001
Enrichment score (ES)
0.2
0.1
0.0
-0.1
Higher in COVID-19 BALF (n=8)
Higher in healthy BALF (n=20)
****
IL10 (log2 RPKM)
4
2
0
Healthy
COVID-19

**Fig. S1. Cellular phenotypes of $CD4^+$ T cells in BALF of patients with COVID-19. a-b**, UMAP representation of scRNAseq showing main clusters of cells from bronchoalveolar lavage fluid (BALF) of patients with COVID-19 and healthy controls (**a**) and dot plot depicting expression of select marker genes for each cluster (**b**). Highlighted in both **a** and **b** are clusters 2 and 12, which represent T lymphocytes. **c**, Dot plot showing expression of select marker genes for clusters of cells depicted in **Fig. 1a**. **d**, GSEA showing genes more highly expressed in bulk RNA-seq of BALF cells obtained from patients (*n*=8) with COVID-19 compared to healthy controls (*n*=20) are enriched in Th1 genes. Bar chart (right) shows the expression of *IL10* mRNA in these samples. Data in **a-b** are from *n*=9 patients with COVID-19 and *n*=4 healthy controls, sourced from GSE145926 and GSE122960. Data in **c** are from the same sources but with *n*=8 patients with COVID-19 and *n*=3 healthy controls (one sample from each was removed due to too low $CD4^+$ T cell numbers). Data in **d** are from *n*=8 patients with COVID-19 and *n*=20 healthy subjects, obtained from HRA000143. **** $p<0.0001$ by Mann-Whitney U-test.

**Fig. S2**

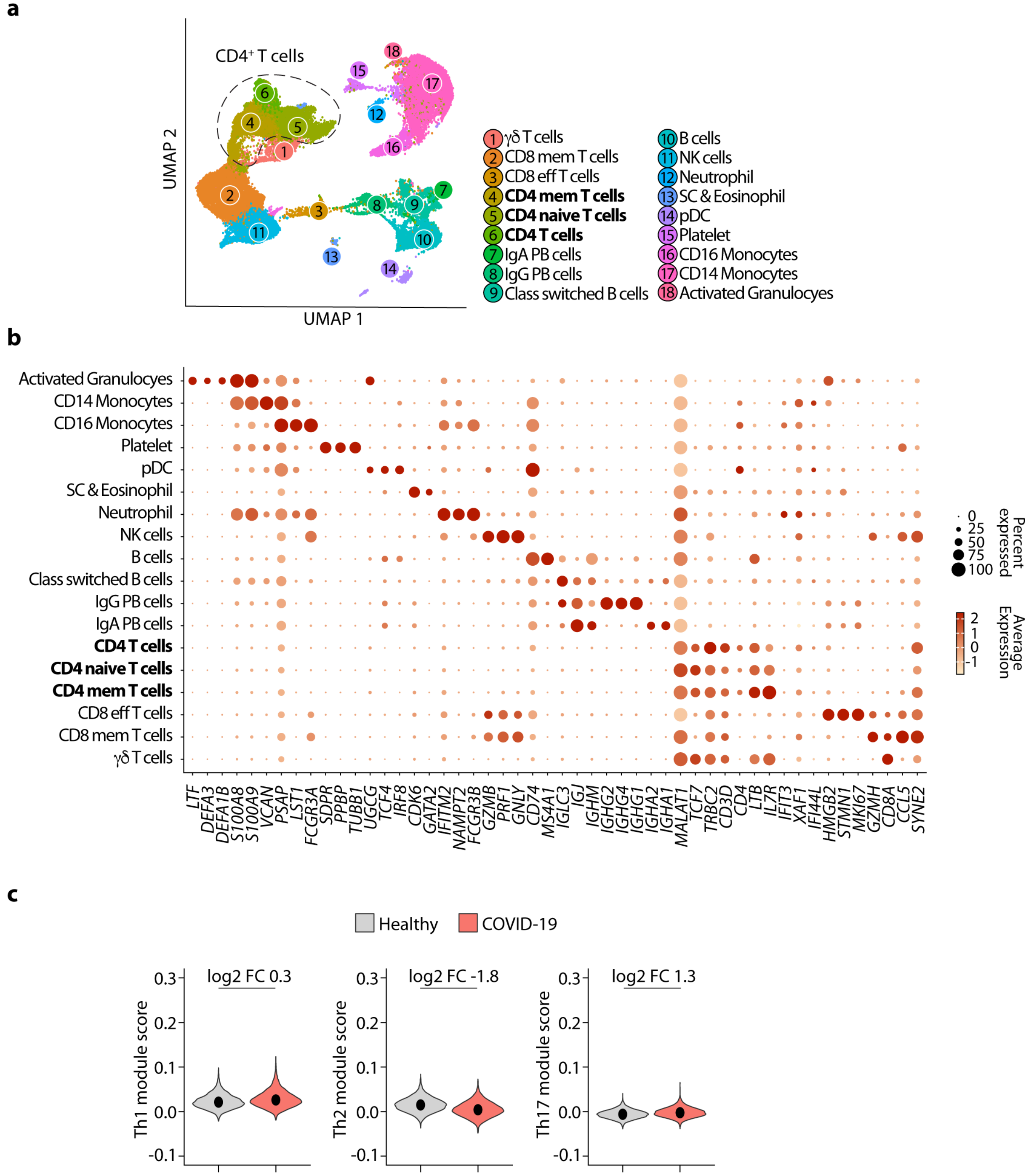


**Fig. S2. Circulating CD4⁺ T cells of patients with COVID-19 are not Th1 biased. a-b**, UMAP representation of scRNAseq showing main clusters of cells from peripheral blood mononuclear cells (PBMC) of patients with COVID-19 and healthy controls (**a**) and dot plot depicting expression of select marker genes for each cluster (**b**). Highlighted in both **a** and **b** are clusters 4, 5 and 6, which represent CD4⁺ T lymphocytes. **c**, Violin plots showing expressions of Th1, Th2 and Th17 genes, respectively, summarized as module scores, in PBMC CD4⁺ T cells of patients with COVID-19 and healthy controls. Data in **a-c** are from *n*=6 patients with COVID-19 and *n*=6 healthy subjects, obtained from GSE150728.

**Fig. S3**

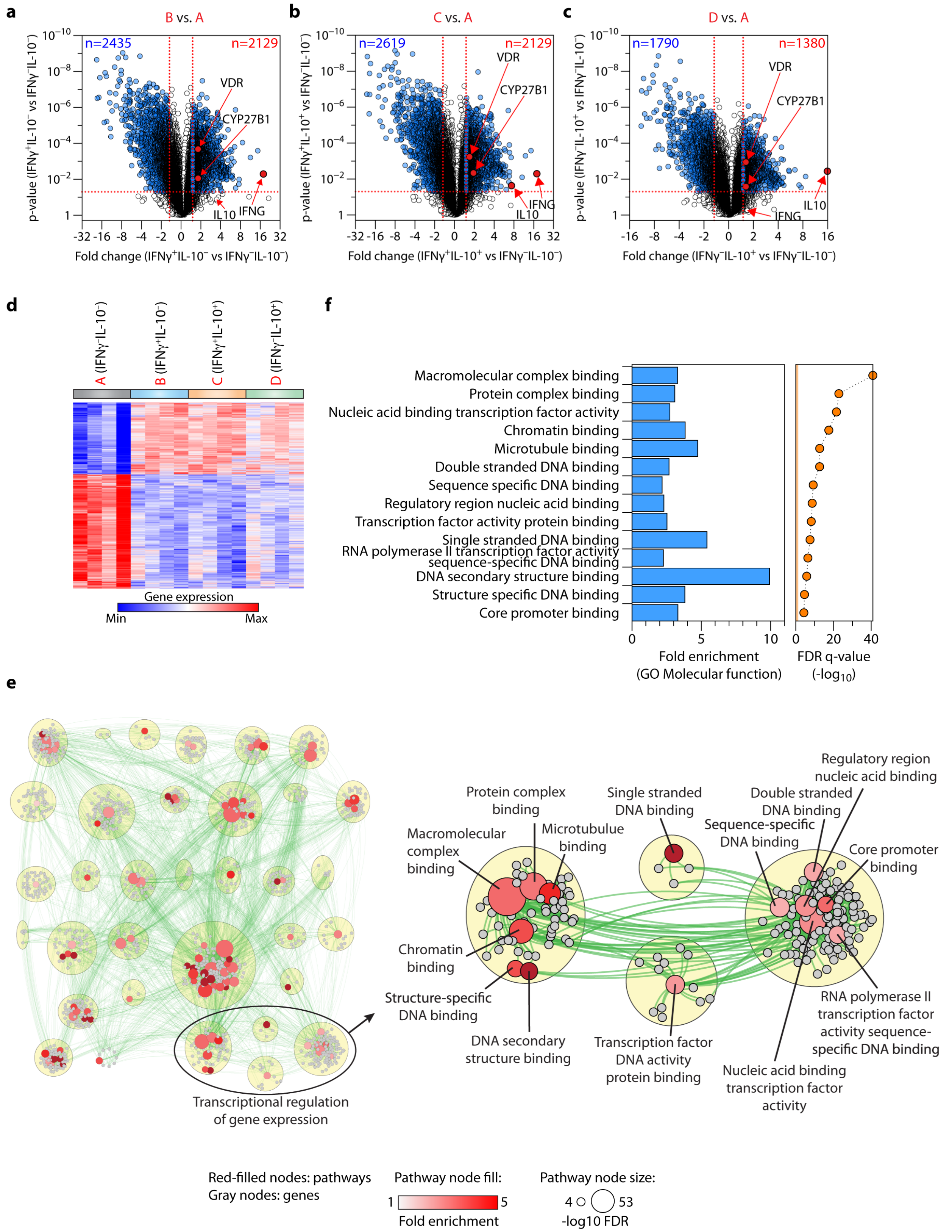

a
B vs. A
n=2435
n=2129
VDR
CYP27B1
IL10
IFNG
b
C vs. A
n=2619
n=2129
VDR
CYP27B1
IFNG
IL10
c
D vs. A
n=1790
n=1380
VDR
CYP27B1
IL10
IFNG
d
A (IFNγ⁻IL-10⁻)
B (IFNγ⁺IL-10⁻)
C (IFNγ⁺IL-10⁺)
D (IFNγ⁻IL-10⁺)
Gene expression
Min
Max
f
Macromolecular complex binding
Protein complex binding
Nucleic acid binding transcription factor activity
Chromatin binding
Microtubule binding
Double stranded DNA binding
Sequence specific DNA binding
Regulatory region nucleic acid binding
Transcription factor activity protein binding
Single stranded DNA binding
RNA polymerase II transcription factor activity sequence-specific DNA binding
DNA secondary structure binding
Structure specific DNA binding
Core promoter binding
Fold enrichment (GO Molecular function)
FDR q-value (-log10)
e
Transcriptional regulation of gene expression
Protein complex binding
Single stranded DNA binding
Regulatory region nucleic acid binding
Double stranded DNA binding
Macromolecular complex binding
Microtubule binding
Sequence-specific DNA binding
Core promoter binding
Chromatin binding
Structure-specific DNA binding
DNA secondary structure binding
Transcription factor DNA activity protein binding
RNA polymerase II transcription factor activity sequence-specific DNA binding
Nucleic acid binding transcription factor activity
Red-filled nodes: pathways
Gray nodes: genes
Pathway node fill:
Fold enrichment
Pathway node size:
-log10 FDR

**Fig. S3. Complement-activated CD4$^+$ T cells are enriched in transcription factors. a-c**, Volcano plots showing differentially expressed genes (DEGs) following activation of CD4$^+$ T cells with α-CD3+α-CD46, comparing IFN-γ$^+$IL-10$^-$ cells (**a**), IFN-γ$^+$IL-10$^+$ cells (**b**) and IFN-γ$^-$IL-10$^+$ cells (**c**) to IFN-γ$^-$IL10$^-$ cells, respectively. DEGs are defined as at least 1.5-fold change in either direction at p-value <0.05. Marked in **a-c** are the *IFNG*, *IL10*, *VDR* and *CYP27B1* genes. **d**, Heatmap showing expression of the 2023 shared DEGs in **a-c**. **e**, ClueGo analysis for molecular function terms in the 2023 DEGs shown in **d** represented as a Cytoscape visualization. Genes are shown in grey, enriched molecular function terms are in red scaled to reflect fold enrichment. Node sizes reflect enrichment significance. Related terms grouped as families within yellow circles. Four such families represent transcriptional regulation of gene expression and are shown in the inset on the right. **f**, The top 14 transcription factor molecular function terms are shown, with associated fold enrichments and FDR q-values. Data in **a-f** are from *n*=4 experiments.

Fig. S4

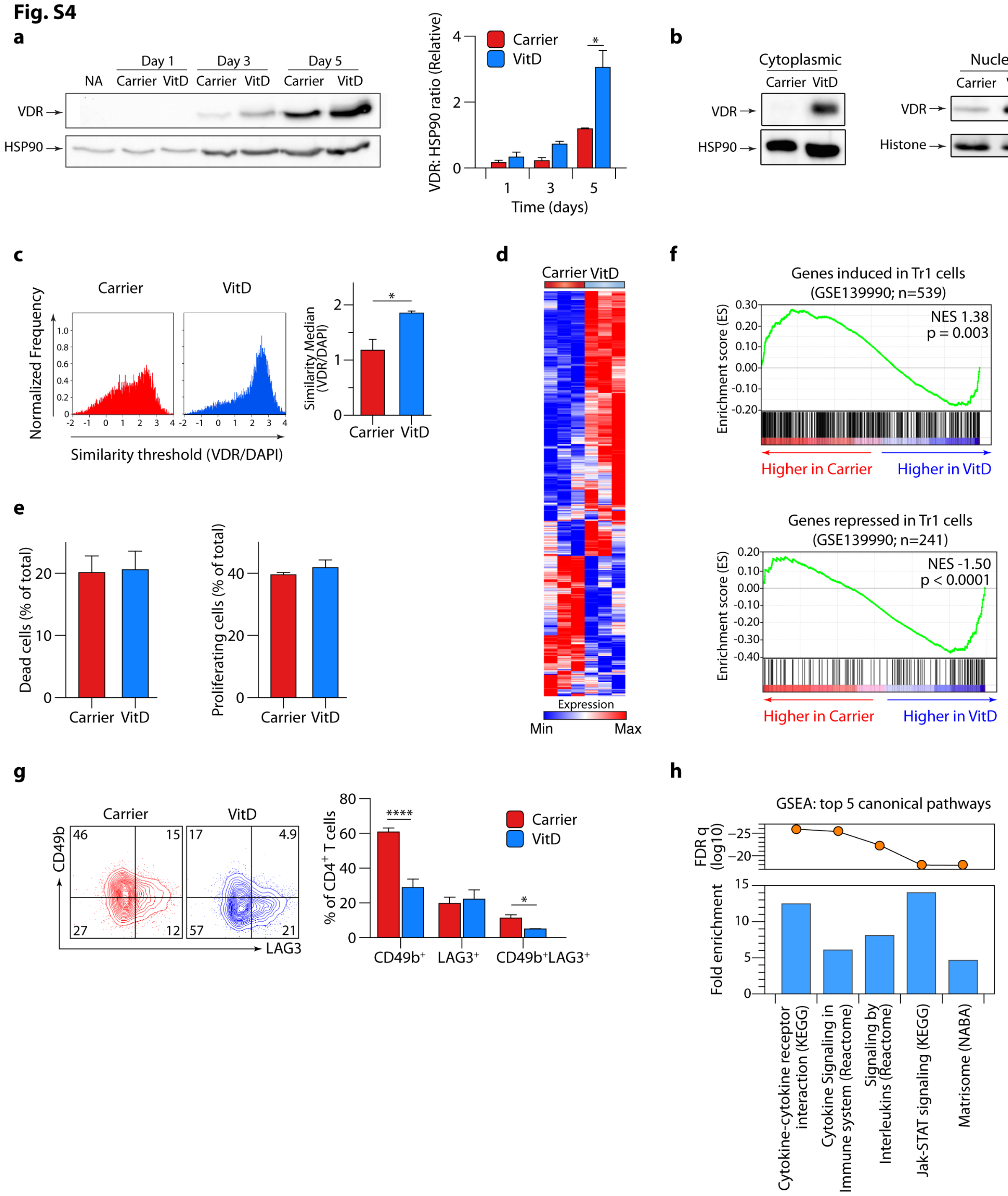


**Fig. S4. Phenotype of Vitamin D treated cells. a**, Western blot (left) and cumulative data (right) for VDR at days 1, day 3 and day 5, with Hsp90 as loading control, in both carrier and VitD treated CD4$^+$ T cells. **b**, Representative immunoblots for VDR and indicated housekeeping proteins in nuclear and cytoplasmic extracts of carrier and VitD treated CD4$^+$ T cells. **c**, Co-localisation of VDR and DAPI in carrier and VitD treated CD4$^+$ T cells, measured on day 2 using ImageStream. Shown are representative frequency histograms indicating overlap between VDR and DAPI in the entire population (left), and cumulative data from *n*=3 independent experiments (right). **d**,

Heatmap showing genes differentially expressed (1.75-fold change in either direction at FDR<0.05) between VitD and carrier treated CD4$^+$ T cells. **e**, Cell death assessed by live/dead stain and proliferation assessed by CFSE dilution in CD4$^+$ T cells treated with carrier or VitD after 3 days of culture. **f**, GSEA showing genes more highly expressed in CD4$^+$ T cells treated with carrier compared to VitD are enriched in Tr1-induced genes (top panel) and genes more highly expressed in CD4$^+$ T cells treated with VitD compared to carrier are enriched in Tr1-repressed genes (bottom panel; curated from GSE139990). **g**, Representative flow cytometry plot showing CD49b and LAG-3 expression in CD4$^+$ T cells, with carrier and VitD treatment (left), and quantification of cumulative data (right). **h**, Top 5 MSigDB canonical pathways enriched in DEGs of VitD vs carrier treated CD4$^+$ T cells (see **Fig. 2a**). Unless indicated, all cells in **Fig. S4** have been activated with α-CD3+α-CD28. Bars represent mean + sem throughout. All experiments have been carried out *n*=3 times. NES = normalized enrichment score. *p<0.05, ****p<0.0001 by 2-way ANOVA (**a**, **c**, **g**) and paired t-test (**c**).

**Fig. S5**

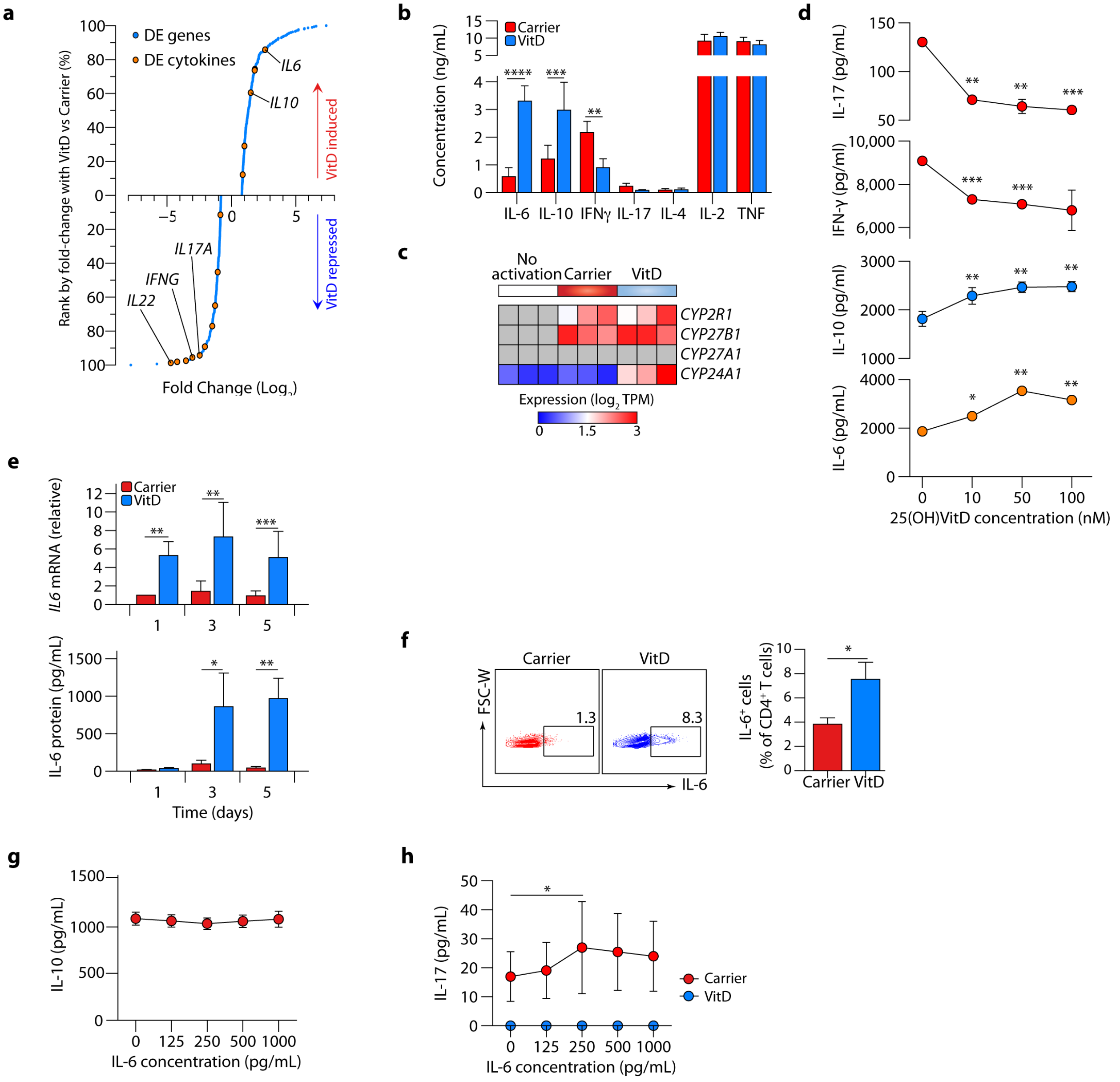


**Fig. S5. Hierarchy of cytokines induced by Vitamin D and modulation of IL-6 effects by Vitamin D. a**, Differentially expressed genes (DEGs) between VitD and carrier treated CD4$^+$ T cells (see **Fig. S4e**) ranked by fold change. Each DEG is marked by a blue dot; each differentially expressed cytokine is marked by an orange dot. Select cytokines have been labelled. **b**, Cytokine concentrations in supernatants of CD4$^+$ T cell cultures after 5 days of treatment with carrier or VitD. **c**, Heatmap showing mRNA expressions (log2 TPM) of the 25-hydroxylase enzymes (CYP2R1 and CYP27A1), the 1α-hydroxylase enzyme (CYP27B1) and the 24-hydroxylase enzyme (CYP24A1), responsible for the two steps of Vitamin D activation and its subsequent inactivation, respectively. Data are from CD4$^+$ T cells activated with α-CD3+α-CD28 and cultured with either carrier or VitD, or left unactivated. **d**, Concentrations of indicated cytokines in culture supernatants of CD4$^+$ T cells treated with escalating doses of 25(OH)VitD for 72h. **e**, *IL6* mRNA, fold change compared to day 1 carrier (above), and IL-6 protein concentration in matched supernatants (below), at days 1, 3 and 5 in carrier and VitD-treated $CD4^+$ T cell cultures. **f**, Representative flow cytometry plot (left) and cumulative data (right) of intracellular IL-6 expression in T cells treated with carrier or VitD (assay carried out on day 3). Cells gated based on lymphocyte gate (forward scatter, side scatter), singlets, live cells and $CD4^+$ cells. **g**, IL-10 concentrations in supernatants of $CD4^+$ T cells cultured in the presence of increasing concentrations of IL-6 for 72 hours. **h**, IL-17 concentrations in supernatants of $CD4^+$ T cells cultured in the presence of increasing concentrations of IL-6, with and without VitD for 72 hours. Unless indicated, all cells in **Fig. S5** have been activated with α-CD3+α-CD28. Cumulative data in **b**, **d-h** depict mean + sem. All experiments have been carried out $n$=3 times. *$p<0.05$, **$p<0.01$, ***$p<0.001$, ****$p<0.0001$ by one-way ANOVA (**d**), two-way ANOVA (**b**, **e**, **h**) and paired t-test (**f**). Statistical comparisons in **d** and **h** compare VitD-treated (**d**) or IL-6-treated (**h**) cells against untreated cells.

**Fig. S6**

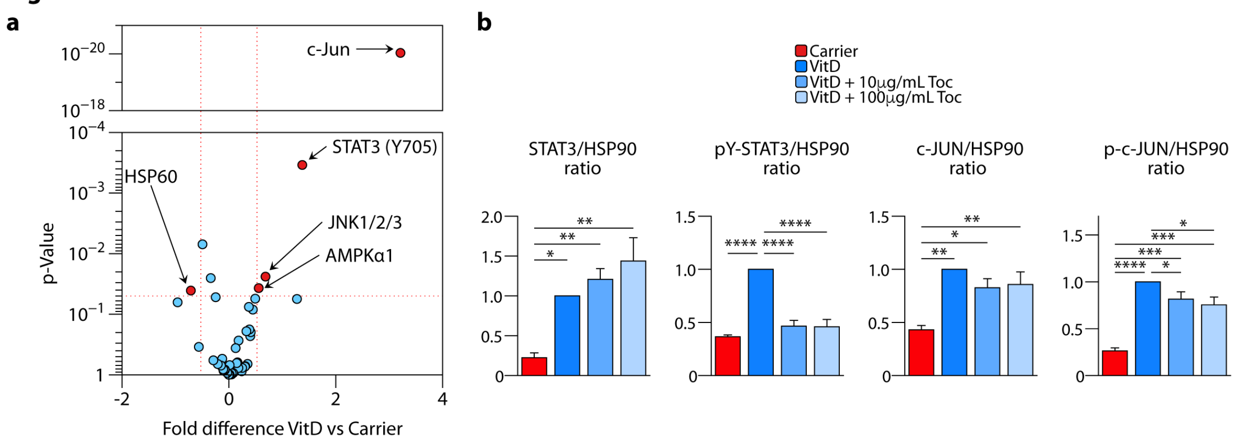


**Fig. S6. Phospho-kinase proteome of Vitamin D-treated cells. a**, Volcano plot representing changes in protein phosphorylation on phospho-kinase array comparing VitD-treated versus carrier treated cells. Data are from $n$=2 independent experiments. Thresholds for significance have been set at 1.2 fold change in phosphorylation in either direction at p-value <0.05. Please also see **Figs. 2f-g**. Marked are phosphoproteins that show significant changes in phosphorylation on VitD treatment. **b**, Quantification of pY-STAT3, STAT3, p-c-JUN, c-JUN and Hsp90 from immunoblots of lysates of $CD4^+$ T cells treated with carrier or VitD with, and without, Tocilizumab (Toc) at the concentrations shown. Please also see **Fig. 2h**. Bars show mean + sem from $n$=3 independent experiments. All cells in **Fig. S6** have been activated with α-CD3+α-CD28. *p<0.05, **p<0.01, ***p<0.001, ****p<0.0001 by one-way ANOVA.

Fig. S7

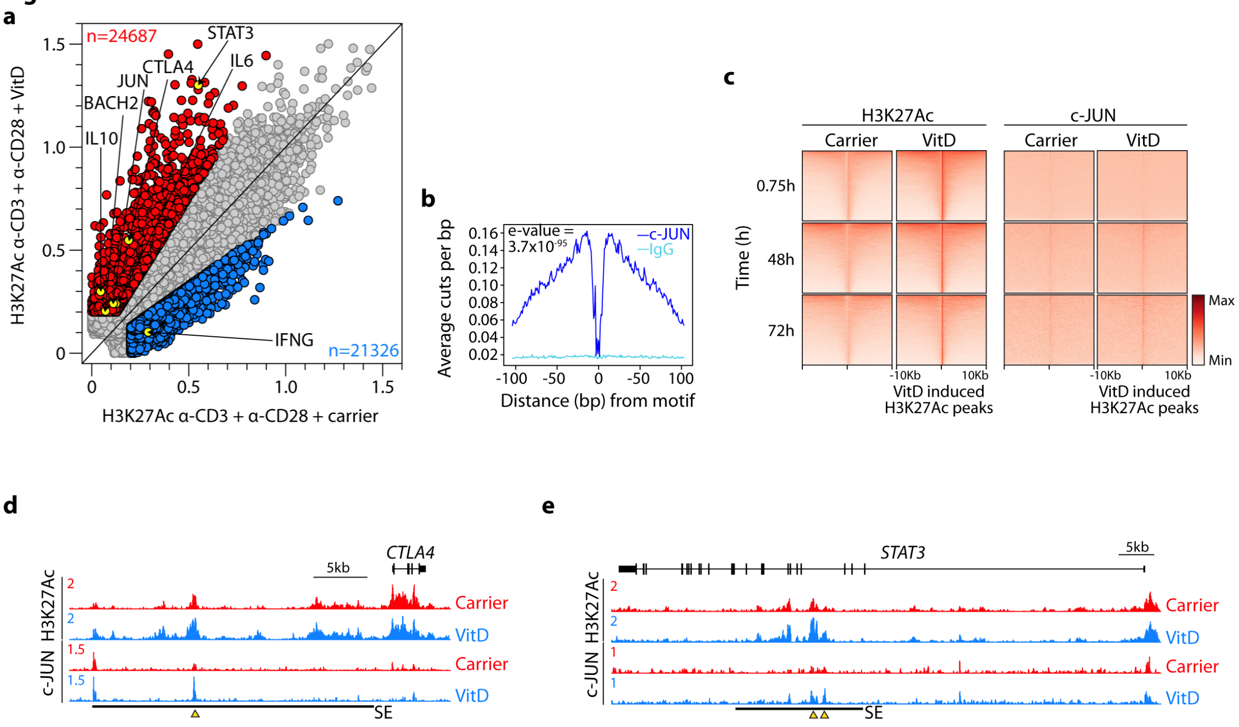


**Fig. S7. Vitamin D alters epigenetic landscapes of CD4$^+$ T cells. a**, Scatterplot showing H3K27Ac CUT&RUN peak intensities 48h after VitD or carrier-treatment of CD4$^+$ T cells. Indicated are VitD-induced peaks (red) and VitD-repressed peaks (blue). Highlighted are select peaks at loci of interest. **b**, histograms showing average cuts per bp in relation to the c-JUN motif in VitD-treated CD4$^+$ T cells. Shown are data from CUT&RUN carried out with IgG (cyan) or c-JUN (dark blue) antibodies. **c**, heatmaps showing H3K27Ac and c-JUN signals at VitD repressed and VitD-induced peaks at the time points indicated. **d-e**, representative genome browser tracks showing H3K27Ac and c-JUN CUT&RUN at the *CTLA4* (**d**) and *STAT3* (**e**) loci following 48hr VitD or carrier treatment of CD4$^+$ T cells. Indicated are the positions of the *CTLA4* and *STAT3* super-enhancers induced by VitD and select c-JUN and/or H3K27Ac peaks (yellow triangles). All cells in **Fig. S7** have been activated with α-CD3+α-CD28.

**Fig. S8**

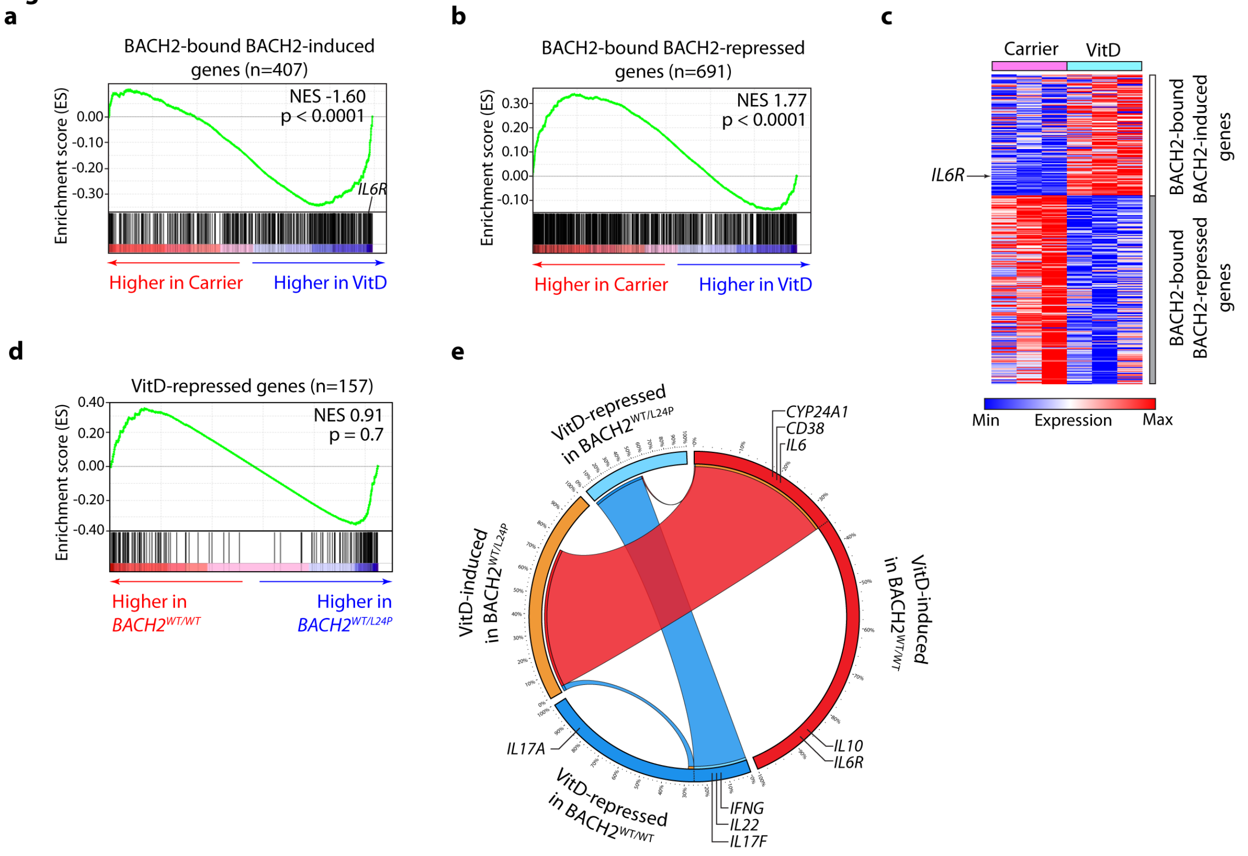


**Fig. S8. A subset of Vitamin D-regulated genes are dependent on BACH2**. **a-c**, GSEA comparing the transcriptomes of carrier and VitD-treated CD4$^+$ T cells against BACH2-bound BACH2-induced genes (**a**) and -repressed genes (**b**). Shown in **c** are the leading edges of the two GSEA enrichment plots in **a-b**. Marked in **a** and **c** is the IL-6 receptor (IL6R). **d**, GSEA comparing enrichment in VitD-repressed genes of the transcriptomes of VitD-treated CD4$^+$ T cells from a patient with haploinsufficiency of BACH2 (BACH2$^{WT/L24P}$) and healthy wild-type controls (BACH2$^{WT/WT}$). **e**, Circos diagram showing VitD-induced and repressed genes in VitD-treated CD4$^+$ T cells from a haplo-insufficient BACH2$^{WT/L24P}$ patient and healthy haplo-sufficient BACH2$^{WT/WT}$ control. Cords join shared genes in patient and control. Indicated are the shared VitD-induced genes (*CYP24A1*, *CD38* and *IL6*) and genes only induced by VitD in the presence of two wild-type copies of BACH2 (*IL10* and *IL6R*). All cells in **Fig. S8** have been activated with α-CD3+α-CD28. NES = normalized enrichment score.

**Fig. S9**

**a**

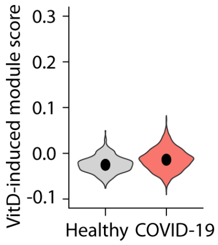


**Fig. S9. Vitamin D-induced genes do not distinguish CD4⁺ BALF T cells of patients with COVID-19 from healthy controls. a**, Violin plots showing expressions of VitD-induced genes, summarized as module scores, of BALF CD4⁺ T cells of patients with COVID-19 and healthy controls. Data are from *n*=8 patients with COVID-19 and *n*=3 healthy controls, sourced from GSE145926 and GSE122960.

**Fig. S10**

**a**

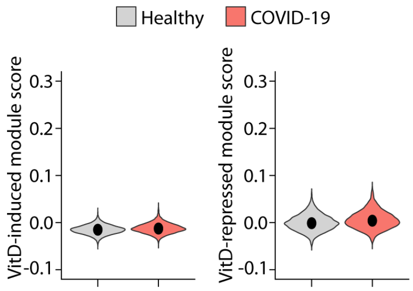


**Fig. S10. Vitamin D-induced or repressed genes are not different between CD4⁺ T cells in peripheral circulation of patients with COVID-19 compared to healthy controls**. **a**, Violin plots showing expressions of VitD-induced and VitD-repressed genes, summarized as module scores, of PBMC CD4⁺ T cells of patients with COVID-19 and healthy controls. Data are from *n*=6 patients with COVID-19 and *n*=6 healthy subjects, obtained from GSE150728.

**Fig. S11**
**a**

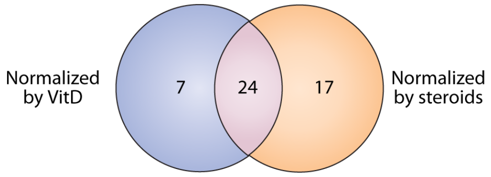


**Fig. S11. VitD and steroid target genes are shared. a**, Venn diagram showing overlap between COVID-induced genes in $CD4^+$ BALF T cells that are predicted to be normalized by VitD treatment versus those that are predicted to be normalized by steroid drugs. Please also see **Table S6**.

**Supplementary Tables**

**Table S1. Differentially expressed genes and enriched pathways between CD4$^+$ BALF T cells of patients with COVID-19 compared to healthy controls**. **a**, List of differentially expressed genes (at least 1.5 change in either direction at adjusted p-value <0.05) between BALF CD4$^+$ T cells of patients with COVID-19 and healthy controls. Data are from GSE145926 and GSE122960. **b**, MSigDB genesets enrichment in DEGs of BALF CD4+ T cells from patients with COVID-19 compared to healthy controls. **c**, Expression of all genes in different Th1 subsets from **Fig. 1f.** (shown are normalized signals from microarray) Data are from GSE119416. **d**, Differentially expressed genes (at least 1.5 change in either direction at p-value <0.05) between different Th1 subsets in **Fig. 1f**. **e**, Enrichment of gene ontology molecular function terms in shared DEGs (intersect of Venn diagram in Fig. 1g) calculated using ClueGO for Cytoscape. Transcription factor terms are highlighted in red. **f**, EnrichR based prediction of transcription factors regulating genes induced in BALF CD4$^+$ T cells (top) and lung biopsies (bottom) of COVID patients compared to healthy controls.

**Table S2. Genes regulated by Vitamin D. a,** Expression of all genes in CD4$^+$ memory T cells activated, as indicated, for 48hr. RPKM values are shown. b, Differentially expressed genes (at least 1.75-fold change in either direction at FDR <0.05) between VitD versus carrier treated cells after 48hr.

**Table S3. Genomic effects of Vitamin D**. **a**, H3K27ac CUT&RUN peaks in carrier and VitD-treated cells at the indicated time points. **b**, Superenhancer and typical enhancer H3K27Ac signals before and after VitD treatment for 48h. **c**, JUN CUT&RUN peaks in carrier and VitD-treated cells after 48h. All coordinates in **a-c** are in hg19.

**Table S4. Response to Vitamin D in BACH2 haplo-sufficient and -insufficient states. a**, Expression (TPM) of all genes in carrier (EtOH) and VitD-treated CD4$^+$ T cells from a haplo-insufficient BACH2$^{WT/L24P}$ patient (Pt.) and healthy haplo-sufficient BACH2$^{WT/WT}$ (HC) control. **b**, Differentially expressed genes (at least 1.75-fold change in either direction at FDR <0.05) between carrier and VitD-treated CD4+ T cells of either a healthy haplo-sufficient BACH2$^{WT/WT}$ (HC) control (left) or haplo-insufficient BACH2$^{WT/L24P}$ patient (Pt) (right).

**Table S5. ROC analyses.** Area under the curve (AUC) statistics for the expression values, summarized as the module score, of all MSigDB hallmark and canonical genesets within CD4$^+$ BALF T cells of patients with COVID-19 compared to healthy controls. Data are from GSE145926 and GSE122960. Pathways have been ranked according to AUC. Marked in red is the Vitamin D repressed geneset.

**Table S6. Drugs predicted by EnrichR to normalize genes induced in CD4$^+$ BALF cells of patients with COVID-19 compared to healthy controls.** Highlighted in red are Vitamin D analogues.

**Table S7. Genesets used in GSEA analysis and module score calculations throughout the paper.**
